# Task-dependent vocal adjustments to optimize biosonar-based information acquisition

**DOI:** 10.1101/2020.08.06.239913

**Authors:** Daniel Lewanzik, Holger R. Goerlitz

**Affiliations:** Acoustic and Functional Ecology, Max Planck Institute for Ornithology, Eberhard-Gwinner-Straße, 82319 Seewiesen, Germany

**Keywords:** active sensing, behavioural flexibility, sensory processing, sensory-motor constraints

## Abstract

Animals need to acquire adequate and sufficient information to guide movements, yet information acquisition and processing is costly. Animals thus face a trade-off between gathering too little and too much information and, accordingly, actively adapt sensory input through motor control. Echolocating animals provide the unique opportunity to study the dynamics of adaptive sensing in naturally behaving animals, since every change in the outgoing echolocation signal directly affects information acquisition and the perception of the dynamic acoustic scene. Here we investigated the flexibility with which bats dynamically adapt information acquisition depending on a task. We recorded the echolocation signals of wild-caught Western barbastelle bats (*Barbastella barbastellus*) while flying through an opening, drinking on the wing, landing on a wall, and capturing prey. We show that the echolocation signal sequences during target approach differed in a task-dependent manner; bats started target approach earlier and increased information update rate more when the task became increasingly difficult, and bats also adjusted dynamics of call duration shortening and peak frequency shifts accordingly. These task-specific differences existed from the onset of object approach, implying that bats plan their sensory-motor program for object approach exclusively based on information received from search call echoes. We provide insights into how echolocating animals deal with the constraints they face when sequentially sampling the world through sound by adjusting acoustic information flow from slow to extremely fast in a highly dynamic manner. Our results further highlight the paramount importance of high behavioural flexibility for acquiring information.

**Summary statement:** Having the right information for a specific job is crucial. Echolocating bats flexibly and independently adjust different call parameters to match the sensory-motor challenges of four different tasks.

## Introduction

Information is the sensory foundation of adaptive behaviors (Dall, Giraldeau, Olsson, McNamara, & Stephens, 2005). Hence, animals constantly integrate a wide range of information to decide on appropriate actions (Gomes et al 2016; Lewanzik, Sundaramurthy, & Goerlitz, 2019; Prat & Yovel, 2020). Acquiring the right information before making decisions is of paramount importance, particularly when decisions have direct fitness consequences, e.g. when failing to evade predators or finding food patches. This universal need for informed decisions is reflected in the evolution of the multitude of senses and their extreme speed and sensitivity (Elemans, Mead, Jakobsen, & Ratcliffe, 2011; Morshedian & Fain, 2015). Yet, animals face the challenge of distinguishing between relevant and irrelevant information (Bates, Simmons, & Zorikov, 2011; Beetz, Hechavarría, & Kössl, 2016), since the central nervous system processing capacity is limited and neural processing is metabolically costly (Laughlin, de Ruyter van Steveninck, & Anderson, 1998; Niven & Laughlin, 2008). Accordingly, selective attention, i.e. the ability to perceptually focus on relevant stimuli while ignoring irrelevant ones, is a crucial feature of cognitive processing (Lavie, 2005) and likely evolved early in evolution (Sridharan, Schwarz, & Knudsen, 2014). In addition, animals apply pre-attentive filters by actively adjusting their primary sensory input, e.g., by controlling the timing (Moss, Chiu & Surlykke 2011; Kurnikova et al 2017; Ladegaard & Madsen 2019) and spatial focus of sensory sampling (Yarbus 1967; Kugler & Wiegrebe, 2017; Geipel et al., 2019; Taub & Yovel, 2020). To study how animals control their primary sensory input, active sensing is an excellent model, as active sensing animals generate and adjust the signal for probing their environment, which we in turn can quantify to investigate the animal’s information acquisition strategy.

Echolocating bats can orient and forage in complete darkness by sound. They control information quantity by changing call rate, and information quality by adjusting various call features, such as duration, amplitude, frequency, and directionality. Despite the large ecological variety of the more than 1200 bat species that echolocate (Wilson & Mittermeier, 2019), every bat needs to perform specific tasks on a daily basis, such as exiting the roost through an opening, drinking from a water body, capturing prey, and landing on surfaces. Each of those tasks constitutes a very different sensory-motor challenge, which bats might solve by adaptively adjusting their biosonar parameters to optimize the rate and features of acquired information. Yet, an adjustment that is aiding navigation during one task might be disadvantageous in another situation. Increasing call rate, for instance, corresponds to an increase of information update rate, which bats commonly perform when approaching objects (Fenton, Jensen, Kalko, & Tyack, 2014; Schnitzler, Moss, & Denzinger, 2003). The most extreme case occurs when approaching flying prey, where aerial-hawking bats emit a characteristic call sequence with extremely high call rates, the so-called feeding (or final / terminal) buzz, right before capture (Griffin, 1958; Kalko & Schnitzler, 1989; Lewanzik et al., 2019; Ratcliffe, Elemans, Jakobsen, & Surlykke, 2013). However, high call rates might be energetically expensive especially for bats calling at high amplitudes (Currie, Boonman, Troxell, Yovel, & Voigt, 2020). High call rates can also cause call-echo ambiguity and thus impair ranging and navigation, when echoes of a call return only after the next call has been emitted (Beetz, Kössl, & Hechavarría, 2019; Moss, Chiu, & Surlykke, 2011). Thus, bats temporally arrange their calls in groups, particularly when a task becomes challenging, so that the longer intervals between groups allow the reception of echoes from distant objects before emitting the next call (Wheeler et al., 2016). In addition, stable call intervals within a call group might aid object localisation and tracking (Moss et al., 2011). Similar trade-offs also exist for other call features: long-duration calls are a means of increasing the emitted energy and, accordingly, the ensonified volume, and thus are often used in open space (Schnitzler & Kalko, 2001). However, long-duration calls increase the probability that echoes reflected off close-by objects overlap in time with the outgoing call (‘forward masking’), which can impair object detection and evaluation (Schnitzler et al., 2003). The same extension of detection distance can be achieved by increasing call amplitude, yet further increasing the amplitude of high-amplitude calls comes at a disproportionately high energetic costs (Currie et al., 2020). Also, high call amplitudes would result in a large number of ‘uninformative’ echoes in cluttered environments, which could distract from relevant echoes and challenge sensory processing (Wahlberg & Surlykke, 2014). Thus, bats usually reduce call amplitude while closing in on targets (Koblitz et al., 2011). Simultaneously, bats often increase bandwidth, i.e. the range of frequencies in their calls, when approaching a target. A broad bandwidth may help to evaluate spectral interference patterns in target echoes but, on the other hand, broad bandwidth calls usually come with decreased detectability since the emitted energy is distributed over a larger frequency range (Wahlberg & Surlykke, 2014).

To date, biosonar adjustments during object approach and prey capture have been well studied in a very limited number of species only. Most of these species are Vespertilionid bats that emit their echolocation calls through the mouth (e.g. Falk, Kasnadi, & Moss, 2015; Hiryu, Hagino, Riquimaroux, & Watanabe, 2007a; Koblitz et al., 2011; Ratcliffe & Dawson, 2003; Russo, Jones, & Arlettaz, 2007; Simmons, Brown, Vargas-Irwin, & Simmons, 2020). Some studies focused on the specialised group of flutter detecting horseshoe bats instead, which emit unique constant frequency calls through the nose (Tian & Schnitzler, 1997; Yamada et al., 2020). Barbastelle bats *Barbastella barbastellus* differ from the majority of other bat species in that they alternately emit two different call types, that are both short and downward frequency modulated. The two call types differ by ∼70° in vertical emission direction (Seibert, Koblitz, Denzinger, & Schnitzler, 2015), call duration (up to 2x), peak frequency (33-34 kHz vs 40-42 kHz), frequency bandwidth (up to 2x), and probably in emission mode (oral vs. nasal emission) (Denzinger, Siemers, Schaub, & Schnitzler, 2001; Seibert et al., 2015). In contrast to most other aerial-hawking species, the calls of barbastelle bats are of low-amplitude that result in very short detection distances, but that allow them to almost exclusively feed on eared moths (Goerlitz, Ter Hofstede, Zeale, Jones, & Holderied, 2010; Lewanzik & Goerlitz, 2018). Despite their low call amplitude, barbastelle bats capture prey on the wing in (semi-)open habitats; however, they usually roost and forage inside forests and thus are used to cluttered conditions (Dietz, von Helversen, & Nill, 2007; Kerth & Melber, 2009; Zeale, Davidson-Watts, & Jones, 2012). Here, we comparatively investigated the biosonar adjustments of barbastelle bats during four object-approach tasks. We hypothesized that bats flexibly adjust their echolocation behaviour in a task-specific way, reflecting the specific echo-acoustic challenges of any given task. We recorded the biosonar of barbastelle bats under controlled conditions in the flight room while bats were flying through an opening, drinking from a pond, landing on a wall, and capturing aerial prey, and we comparatively quantified their task-dependent information acquisition behaviour.

## Materials and methods

### Capture and husbandry

In August 2018, we captured three adult male barbastelle bats *Barbastella barbastellus* using mistnets (16 × 16 mm^2^ mesh size; Ecotone, Gdynia, Poland) at emergence from a roost in eastern Bavaria (Germany) and transferred them to the husbandry facilities of the Max Planck Institute for Ornithology on the night of capture. We kept the bats for about three weeks in an aviary (1.2 × 2.4 × 2.0 m^3^, W × L × H) before releasing them at the site of capture. We provided *ad libitum* access to fresh water at all times. Live moths and/or mealworms (larvae of *Tenebrio molitor*) were provided *ad libitum* during training and experiment in a flight room. If bats had not reached their initial body mass after foraging in the flight room, we additionally hand-fed mealworms before releasing bats back into the aviary.

### Experimental setup

Experiments were conducted in an echo-attenuated flight room. We divided the flight room into a ‘retreat compartment’ (2.7 × 3.4 × 3.0 m^3^, L × W × H) and a ‘foraging compartment’ (5.6 x 3.4 × 3.0 m^3^, Fig. 1) by a combination of a wooden wall at the bottom and a thick felt curtain on top. Bats could fly between compartments through a rectangular opening (size: 1.3 x 0.5 m^2^, W x H) in the wooden wall, whose centre was at 0.75 m above ground and 1.05 m from the closer side wall. The foraging compartment contained a large pool (1.4 x 2.3 m^2^) to drink and a movable stand to offer tethered mealworms (1.92 m above ground). During half (17/35) of the prey capture trials, we ran a “flutter simulator” about 10 cm next to the mealworm. The flutter simulator, a small plastic propeller of the size and shape of a single wing of *Noctua pronuba*, one of the barbastelle’s main prey insects (Goerlitz et al., 2010), imitated the wing beat of a flying *Noctua pronuba* (ca. 43 Hz; Waters & Jones, 1994). During the other half of prey capture trials, the propeller engine was running as well but the wing was detached. Since we did not find any effect of the propeller (see supplementary information), we pooled the data for all other analyses. We positioned four microphones (CM16/CMPA, Avisoft Bioacoustics, Glienicke, Germany) to record the bats’ echolocation behaviour at various positions: two microphones at both short ends of the pool at about 10 cm above ground pointing towards the centre of the pool; one microphone in a corner of the foraging compartment close to the site at which bats landed most often during training, pointing towards the bats’ common approach direction; and one microphone at the top of the mealworm stand pointing towards the tethered mealworm. A fifth microphone (FG-O, Avisoft Bioacoustics) was placed at the side of the opening at 98 cm above ground, pointing towards the bats approach direction from the foraging compartment. Calls on CM16 microphones were recorded with custom-written Matlab code (MathWorks, Natick, MA, USA), SoundMexPro software (HörTech gGmbH, Oldenburg, Germany), a Scarlett OctoPre preamplifier (Focusrite Plc, High Wycombe, UK), and a soundcard (Fireface 802, RME, Haimhausen, Germany; 192 kHz sampling rate, 16-bit resolution). Calls on the FG-O microphone were recorded at 500 kHz sampling rate using UltraSoundGate 416H (Avisoft Bioacoustics) and Avisoft Recorder software. We video recorded the bats’ drinking, foraging, and compartment switching behaviour with an infra-red sensitive camera (HDR-CX560, Sony Corporation, Tokyo, Japan). We placed infrared lights behind the opening inside the retreat compartment perpendicular to the bats’ flight trajectories, such that bats got illuminated in the moment they passed the opening. Further infrared lights illuminated the pond and the tethered mealworm. Video recordings were also used to distinguish successful prey captures from unsuccessful prey capture attempts, both of which ended with a feeding buzz.

**Fig 1:**
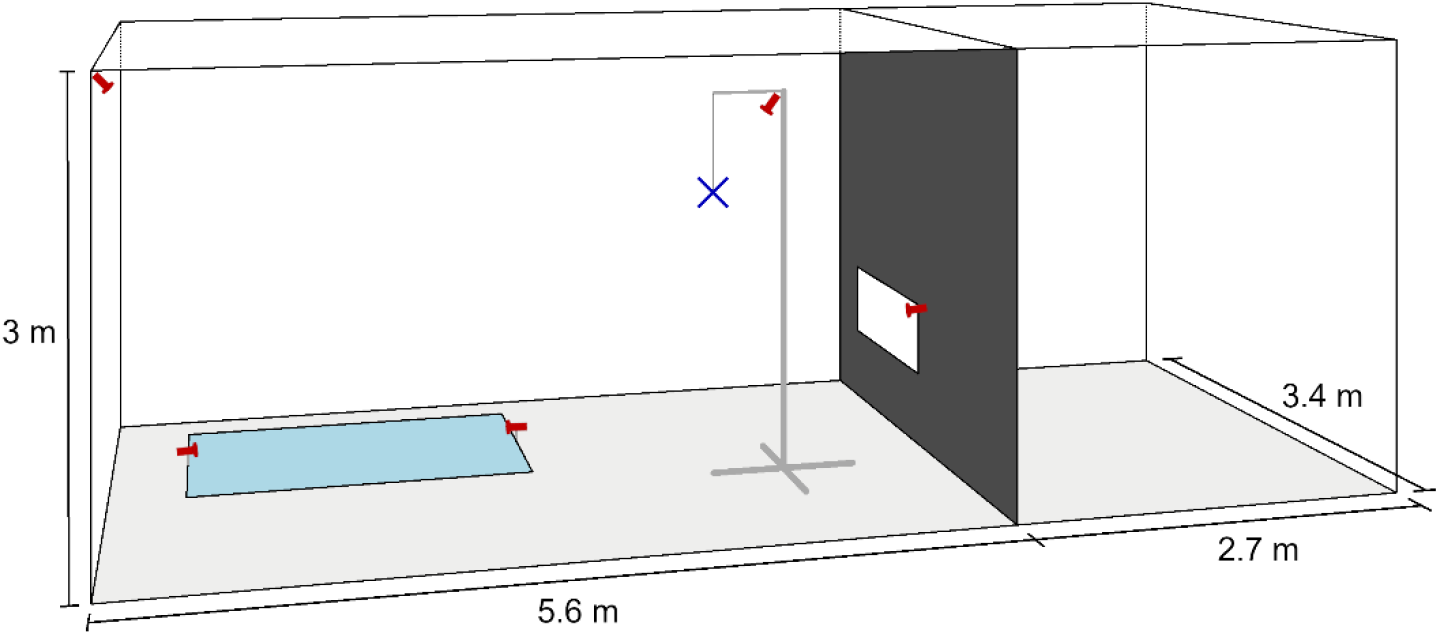
Schematic of the flight room. The flight room was divided by a wall into a foraging compartment (left) and retreat compartment (right). The foraging compartment contained a pool to drink and a single mealworm (blue cross) on a movable stand. The five microphones are shown in red.

**Fig 2:**
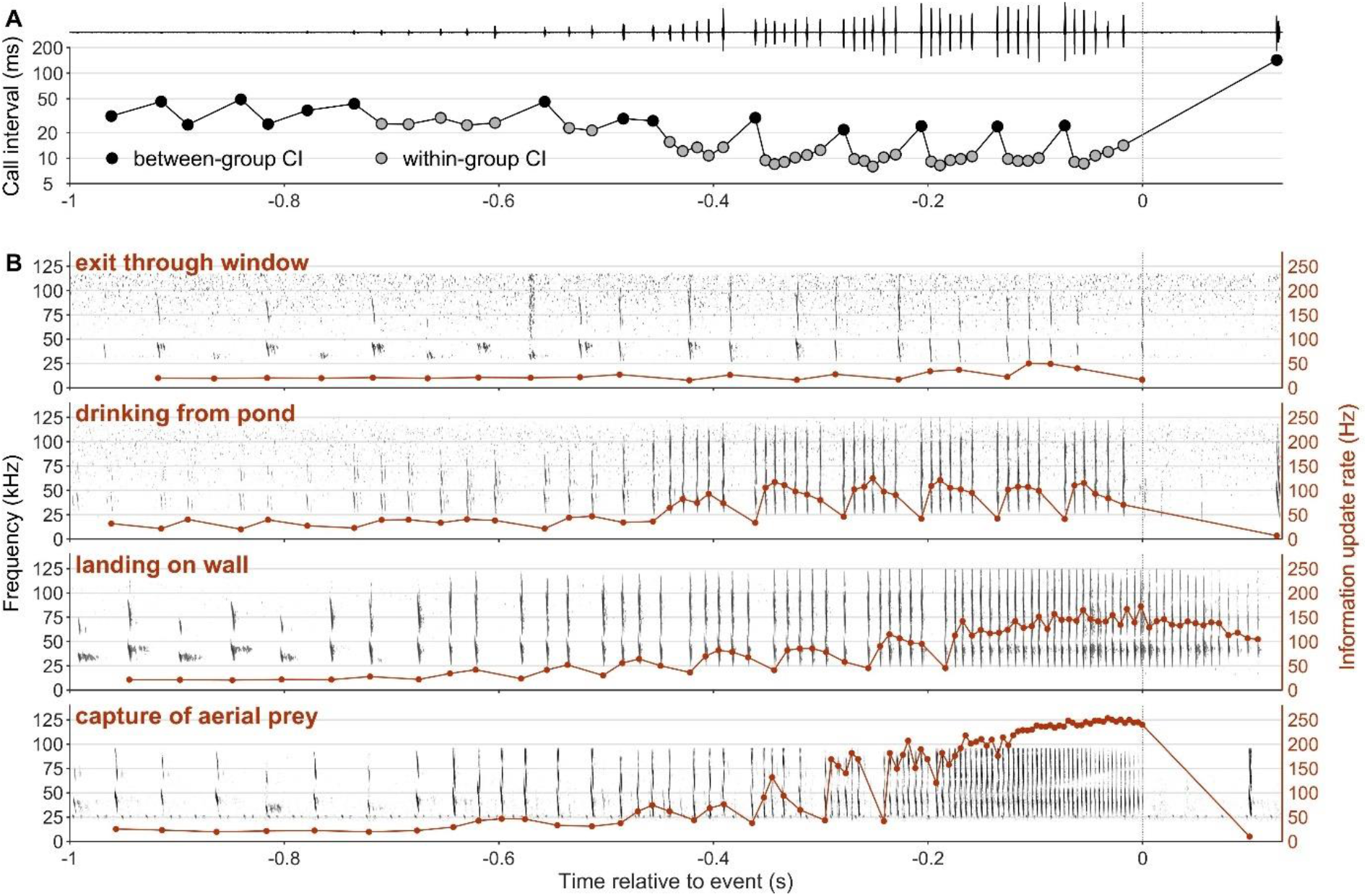
Exemplary object approach call sequences and associated change in call timing. A) Waveform of an exemplary call sequence during a drinking approach, and the call intervals (CIs), separated in between-group and within-group CIs. B) Spectrograms of exemplary call sequences during all four object approach tasks The information update (= call repetition) rate is overlaid and increases during object approach in a task-dependent way.

### Experimental procedure

In the first two nights after capture, we let all bats fly together inside the flight room with multiple tethered live moths and mealworms offered as prey inside the foraging compartment. During the following days, we let bats fly singly in the flight room in random order. Once bats learned to capture tethered prey, we progressively reduced the number of prey items until only two fixed positions were enough to feed the bats. We then started to record the bat when it was drinking or landing while continuously replacing removed prey at the two fixed positions to feed the bats. After multiple nights with two fixed prey locations, we started to only offer a single tethered mealworm at a time at one position (the movable stand), and started to record the bat when attacking this prey item and also when flying through the wall opening from the foraging compartment into the retreat compartment (‘exit’). After bats had captured the mealworm, they usually flew into the retreat compartment. We then closed the opening, replaced the mealworm and moved its stand to one of six positions in random order. After re-opening the opening between the compartments, bats usually flew immediately into the foraging compartment. Once a bat stopped foraging and began grooming itself, we captured it using a hand net, weighed it, hand-fed it more mealworms if necessary, brought it back into the aviary, and released the next individual into the flight room. Every individual could forage in the flight room every night.

### Sound analysis

We analysed the echolocation behaviour of the bats when approaching the pond to drink, the wall to land, the wall opening to exit the foraging compartment, and the prey to capture. Particularly, we compared the echolocation behaviour during the approach phase between tasks since - in contrast to search calls - bats lock their approach calls onto the target (Koblitz et al., 2011; Surlykke, Ghose, & Moss, 2009), indicating the bats’ intention to execute a task and minimizing variation in measured call parameters due to variation in sonar beam direction. Thus, parameters of approach but not of search phase calls are likely adjusted to a given task.

In total, we analysed the sound recordings (wav-file) of 98 object-approach flights using SASLabPro software (Avisoft Bioacoustics). Since landing and drinking were relatively rare events, we analysed all available recordings of landings (N=15) and chose similar numbers of drinking / drinking attempt recordings (N=18 and 15, respectively; at least 5 per individual) with a good signal-to-noise ratio (as evaluated by inspecting the waveform) from across experimental days. For exit, capture and capture attempt flights, we only analysed recordings from the last two nights, i.e. on nights 22 and 23 after capture, when the bats were familiar with the setup and reliably foraged on the mealworm (N=15, 16, and 19 recordings with at least 5 recordings per individual and task). We first highpass-filtered the recordings at 20 kHz (Hamming window, 1024 taps) and downsampled those recordings that were initially recorded with a higher sampling rate to 250 kHz before setting a fixed threshold (−68 dB FS, i.e. rel. to full scale) to detect bat calls above the noise floor. We then manually removed echoes whose amplitude was above detection threshold. For all calls above threshold (N = 5778), peak frequency (from the spectrogram-derived mean spectrum), call duration (as defined by the call detection threshold), and call interval were calculated automatically. However, we subsequently discarded frequency information from 43 calls since these calls were ‘clipped’, i.e. their amplitude exceeded the maximum amplitude that the wav-file could encode. We defined a call’s peak frequency to be within the first or second harmonic if it was below or above 55 kHz, respectively (compare Fig. S1). We then manually determined ‘call type’ (search type I & type II, approach, buzz) of a large subset of calls (3930 calls = 68%) representing all individuals. In barbastelle bats, type I and type II search calls and approach/buzz calls are easily distinguishable by their frequency-time structure (Denzinger et al., 2001; Lewanzik & Goerlitz, 2018; Seibert et al., 2015). To avoid analysing longer call intervals than used by the bats, we added the approximate emission time of 228 calls manually, which were visible in the spectrogram, but below our call detection threshold. Other parameters than the call interval of these calls were not considered for analysis, given that their amplitude was below detection threshold. We used distributions of inter-call-intervals and peak frequency of the manually determined calls to identify ‚call type’ of all remaining calls as follows: We defined calls as search calls when both preceding and following inter-call-interval were ≥36 ms and as approach/buzz calls otherwise. Among search calls we distinguished between type I and type II search calls based on peak frequency (type I: peak frequency either ≤37 kHz or between 55 kHz and 66 kHz; type II otherwise). This might have introduced a few mis-classifications, yet overlap between peak frequency distributions of type I and II search calls is low (Fig. S1) and, importantly, our analyses are not based on search but on approach/buzz calls. To provide values for comparison, we calculated mean and standard deviation of type I and type II search call durations. Since search calls are not target-oriented and thus not necessarily emitted towards the microphone, they were often recorded with low amplitudes, which could bias call duration measurements. To minimise this bias, we used only the 20 % loudest search calls when calculating duration metrics. Among approach/buzz calls, we classified calls as ‘buzz call’ if either the preceding or following inter-call-interval was ≤6 ms; otherwise, they were classified as approach calls. Three individual buzz calls from the same individual were manually re-classified and termed upward-modulated (UM-type) calls since their time-frequency structure differs from all common call types (see results and Fig. S2).

Since bats interspersed approach calls within search call sequences, we defined the onset of the final approach to the object as that approach call after which only approach and buzz but no search calls followed until the ‘event’ as defined below (with the exception of two ‘exit trajectories’ in which bats used a search call already shortly before exiting through the window; Fig. 3 & Fig. S3). The onset of the final buzz was defined accordingly, i.e. as that buzz call after which bats only emitted buzz calls. The ‘event’ in the four object approach tasks was flying through the window, touching the water surface, landing on the wall, and catching the prey, and we used the following criteria to estimate the timing of these events. For both successful prey captures and unsuccessful prey capture attempts (distinguished via video recordings), we defined the event time as the time of the last buzz call. We distinguished successful drinking events from unsuccessful drinking attempts by the splashing sound produced when the bat touched the water, which we used as event time in actual drinking events (Fig. 3), but which was absent in unsuccessful drinking attempts. In the latter case, we defined the event time as the time of the last approach call, which approximates the behaviour in successful drinking events. For landing events, we used the sound of wall contact as event time, which was not always unambiguously visible in the spectrogram but always audible when listening to the sound file. For exit events, i.e. when flying from the foraging into the retreat compartment, we used the last call above call detection threshold as event time (because bats passed a wooden room divider, their calls were not recorded anymore once they flew through the opening). We termed the duration between event time and first post-event call ‘post-event pause’.

**Fig 3:**
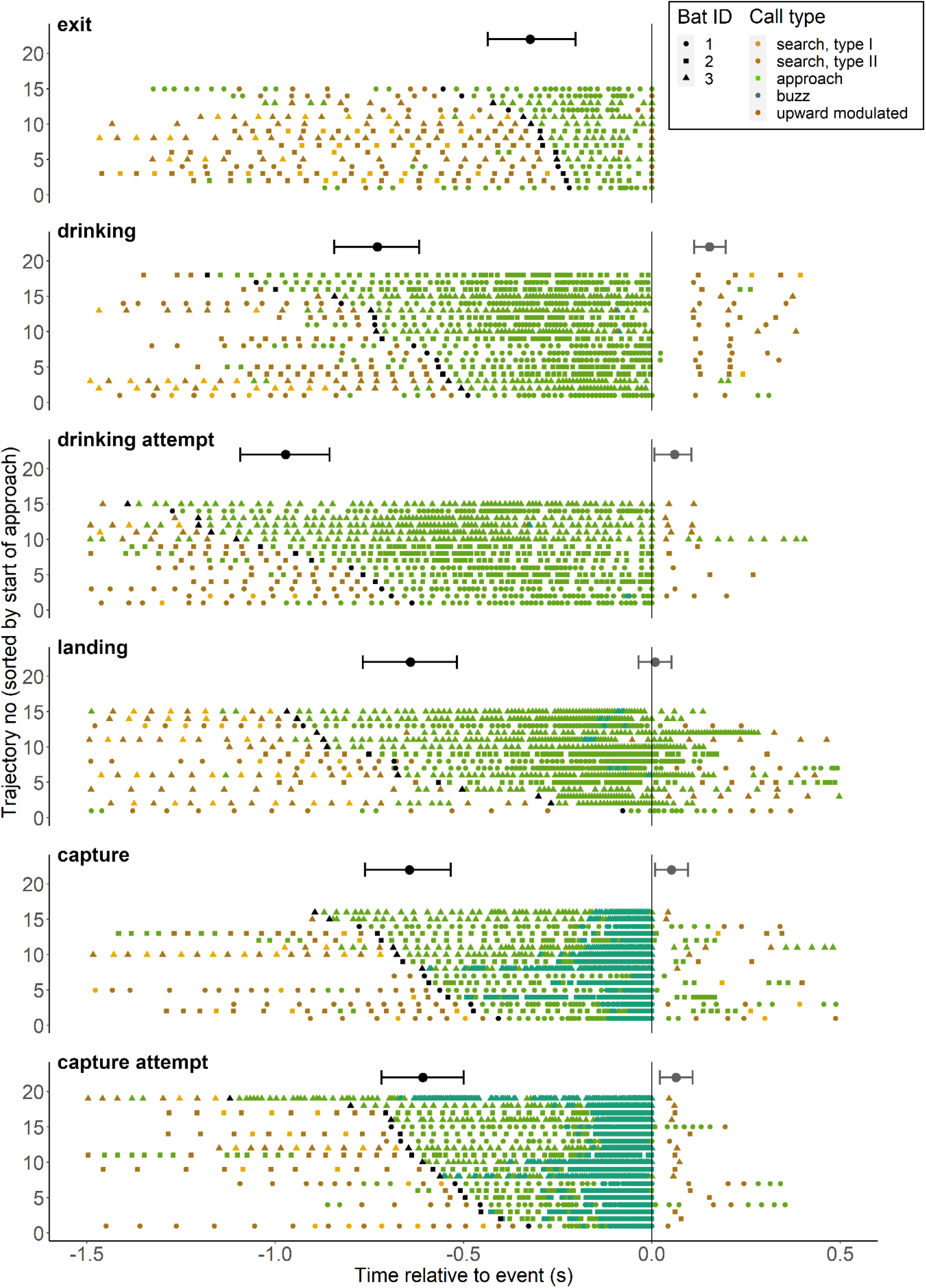
Dynamic emission of different call types during object approach. The start of the final object approach is indicated by black symbols for each trajectory. Horizontal error bars indicate mean posterior predictions and credible intervals from model simulations for the start of the final object approach (black) and for the timing of the first post-event call (grey).

### Statistical analysis

We used the statistical freeware R version 3.6.2 (R Core Team, 2019) to analyse durations of approach phase and post-event pause as well as temporal dynamics of call interval, duration, and peak frequency, and to compare these parameters between tasks. We analysed only the calls during the bats’ final approach to the object, i.e., 2976 approach and 1503 buzz calls, but not search calls. Since bats grouped their calls, we analysed inter-call-interval separately for the shorter within-group intervals and the longer between-group intervals. We defined the first intra-group interval of any call group as the interval that was at least 30% shorter than the preceding call interval or that was absolutely shorter than 10 ms. Subsequent call intervals were considered intra-group intervals as long as they differed maximally by 20% or 5 ms compared to the mean intra-group interval of the current call group (Fig. 2A & Fig. S3).

We fit generalised linear mixed-effects models with a Gaussian error distribution to our response variables ‘onset of final approach’ (rel. to the event time), ‘post event pause’, ‘call duration’, and ‘inter-call interval’ (both within-group and between-group) using the lmer function of the lme4 package (D. Bates, Maechler, Bolker, & Walker, 2015). ‘Onset of final object approach’ and ‘post event pause’ were modelled as a function of the fixed effects ‘event type’ (categorical with six levels: drinking, drinking attempt, exit, landing, prey capture, and unsuccessful prey capture attempt) and – to account for repeated measures – the random effect ‘bat ID’ (categorical with three levels). ‘Call duration’ and ‘inter-call interval’ were modelled as a function of the fixed effects ‘time before event’ (continuous), ‘event type’, and the interaction between the two. Further, we included a 3^rd^ order polynomial of ‘time before event’ as well as its interactions with ‘event type’ as predictors in the models since ‘time before event’ had non-linear effects on these response variables. To prevent autocorrelation, we also included ‘call duration of previous call’ or ‘call interval of previous call’ (continuous) as predictor in these models. Random effects included were ‘trajectory no’ (categorical with 98 levels) to account for potential dependency among calls within the same trajectory, and ‘bat ID’.

To model changes in the proportion of calls with peak frequency in the 2^nd^ harmonic over time, we first calculated the number of calls per 0.3 s time bins with peak frequency in the 1^st^ and 2^nd^ harmonic, respectively (i.e. < or >= 55 kHz; compare Fig. S1). We then fitted a generalised linear mixed model with binomial error distribution to the response variable ‘proportion of calls with peak frequency in 2^nd^ harmonic’ (weighed by the total number of calls per bin) using the glmer function of the lme4 package; including the fixed effects ‘time bin’ (modelled as continuous variable) as a 3^rd^ order polynomial, ‘event type’, the interaction thereof, and the random effects ‘trajectory no’ and ‘bat ID’. In contrast to all other models, we here included data of the entire object approach call sequence (i.e. including also search and approach calls before the beginning of the final approach phase) because binning the data substantially reduced sample size and would not allow modelling polynomial effects otherwise.

All continuous predictors except ‘time bin’ were scaled by their standard deviation and (with the exception of ‘time before event’ and ‘time bin’) centred on their mean. Model assumptions were verified by plotting residuals versus fitted values and by inspecting QQ-plots of the model residuals and of the random effects. We assessed residuals for temporal dependency by plotting estimates of the autocorrelation function using R’s acf function.

We used the sim function of the R-package arm (Gelman & Su, 2016) to simulate the values of the posterior distributions of the model parameters (1,000 simulations) and to obtain posterior mean parameter estimates. We extracted 95% credible intervals (CIs) around the posterior mean estimates and considered support for an effect as strong if zero was not included within the 95% CI and as low if overlap with zero was small, respectively. If 95% CIs were centred on zero, we considered this as strong support for the absence of an effect. We also used the sim function to obtain mean predictions of response variables and 95% CIs around these predictions for various combinations of predictors for plotting purposes. Since we included ‘previous call duration’ and ‘previous call interval’ as predictors in models for call duration and call intervals, respectively, but values of these predictors change over the course of the approach, we used a ‘moving average’ approach to calculate predictions of the other response variables (instead of using the mean value; size of the moving window: 0.3 s and 0.05 s for call times before/after -0.3 s).

## Results

We investigated how flexibly bats adjust their echolocation to control their primary sensory information when executing various tasks, i.e. when approaching an opening to fly through, a pond to drink, a wall to land on, and prey to capture. In search flight, i.e. before initiating object approach, bats usually emitted an alternating pattern of type 1 / type 2 search calls (Fig. 2, Fig. 3 & Fig. S3), which is typical for barbastelle bats. Peak frequencies of the first harmonic of type I and type II search calls were 33.9 ± 1.6 kHz (mean ± sd; N = 126) and 41.5 ± 2.2 kHz (N = 740), respectively (Fig. S1). At times, search calls were interspersed with more broadband approach-like calls. Subsequently, bats stopped emitting search calls altogether and switched to approach and buzz calls only (Fig. 2, Fig. 3 & Fig. S3) with first harmonic peak frequencies of 41.3 ± 3.4 kHz (mean ± sd; N = 2656) and 40.2 ± 2.4 kHz (N = 1278), respectively (Fig. S1). In trajectories, in which we recorded search calls, the last search call was mostly (in 72/82 trajectories: 88%) a type II search call. Both the similar peak frequency and the time-frequency structure suggest that approach and buzz calls are more similar to type 2 than type 1 search calls, likely sharing the same mechanism of vocal production and (nasal) emission. Besides these common call types, one bat also emitted an unusual, upward modulated call in three instances before drinking or attempting to drink, that we termed UM-type call (Fig. S2). We speculate that these UM-type calls were unintentionally emitted when the bat widely opened the mouth in preparation to drink and did not precisely control the timing of exhalation and sound production, respectively.

We defined the beginning of the approach phase as the time of the first approach call after which bats did not emit any further search calls. Approach phase duration differed between tasks, indicating different needs for sensory information. During drinking, landing and prey capture, the bats started the object approach at about 0.6-0.7 s before the event (Fig. 3 & Fig. S3, tables S3/S4). In contrast, when exiting through the opening, the bats started object approach later at about 0.3 s before the event, resulting in a ca. 50% shorter approach duration. In drinking attempts (in contrast to actual successful drinking events), approach duration was longer than in all other tasks (Fig. 3 & Fig. S3, table S5), likely due to continued emission of approach calls while the bats continued trying to drink.

Next, we analysed the dynamic changes and task-dependent differences of temporal and spectral call parameters during the final object approach (i.e., only approach and buzz calls). We will present results for call intervals, call durations, and call peak frequencies. Neither of these, nor the duration of approach phase, buzz, or post-event pause differed between capture trials with and without the flutter simulator (tables S1/S2).

### Temporal patterning of information acquisition

In all tasks, bats decreased their call intervals during the object approach (Fig. 4), thereby increasing the amount and update rate of sensory information (Fig. 2B). Bats also grouped their calls into groups, with shorter within-group call intervals (mean ± sd: 11.2 ± 11.7 ms) and longer between-group call intervals (45.9 ± 35.0 ms; Fig. 2 & Fig. S3). Beyond those common temporal patterns, the dynamics of both within- and between-group call intervals differed between tasks (Fig. 4, tables S3-S5): bats initially reduced both call intervals during the approach phase, but then stopped reducing (landing, drinking) or even increased call intervals towards the end of the approach (exiting, drinking attempts). In contrast, only during prey capture (attempts), bats kept on reducing call intervals until the event. In addition, the bats started their approach already with shorter call intervals during prey capture (attempts) than during any other task, reducing them further to the shortest call intervals right before capturing prey (min: 3.5 ms). Also, terminal buzzes, i.e. sustained sequences of buzz calls (call interval < 6 ms) were only emitted during prey capture (attempt) tasks (Fig. 3, Fig. 4b, Fig. S3). The onset of the buzz was at 169 ± 76 ms (mean ± sd) before capture and did not differ between successful and unsuccessful prey capture trials (tables S3/S4). Note, however, that there was no clear distinction in call interval between approach and buzz phases, but rather a gradual transition.

**Fig 4:**
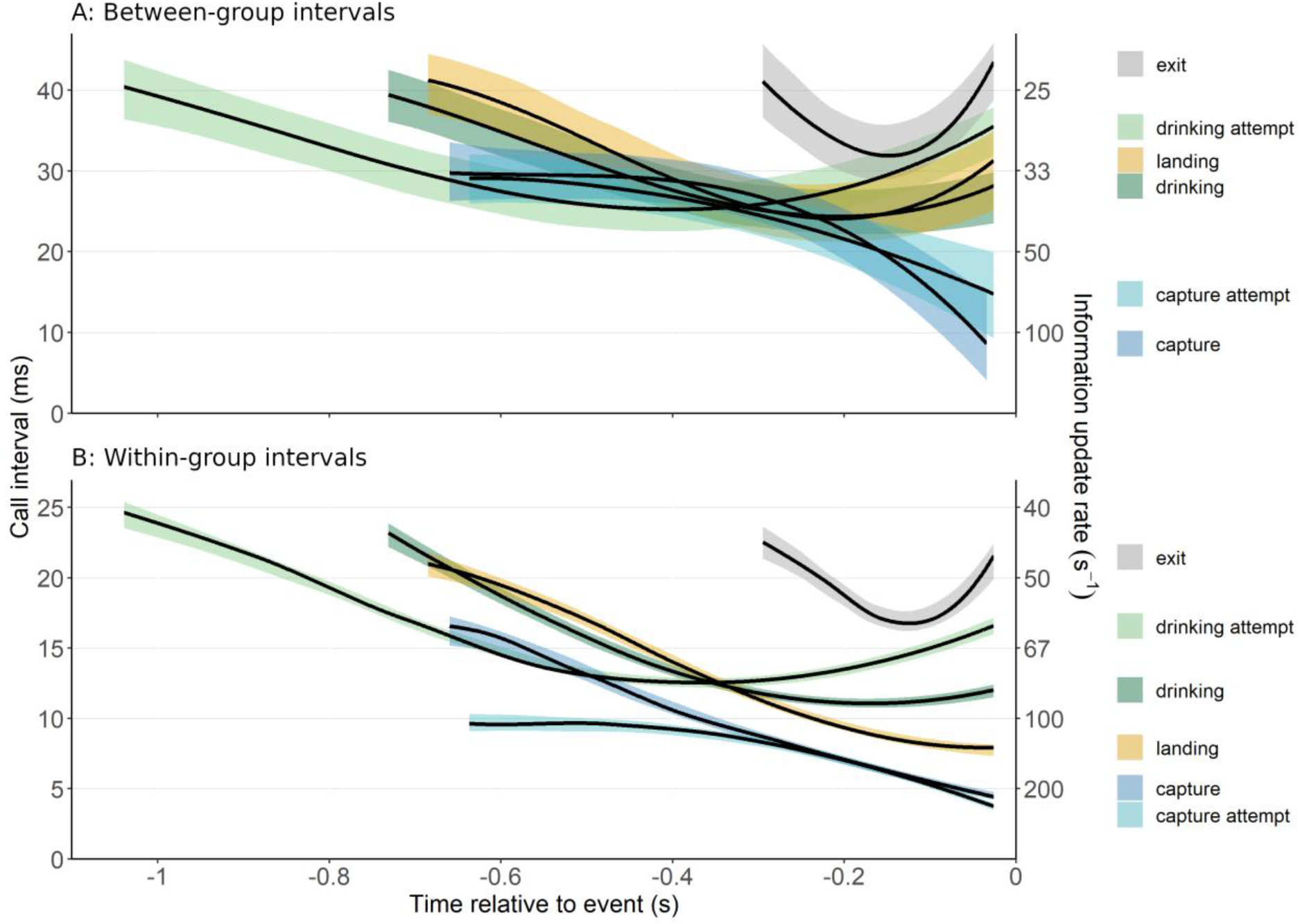
Dynamics of call interval (left y-axis) and corresponding information update rate (right y-axis) during object approach. Call intervals are separated into A) between-group and B) within-group call intervals. Depicted are model predictions (solid lines) and 95% credible intervals (coloured bands). Models are based on N=1029 between-group and N=3254 within-group call intervals, respectively. Predictions are made starting from the event-specific median start time of the final (object) approach phase.

As a result of these different dynamics, update rates right before the event were lowest when approaching the opening to exit the foraging compartment, intermediate when approaching the pool to drink, high before landing, and highest (up to max. 286 s^-1^) before capturing prey (Fig. 2B, Fig. 4). With the exception of exit and landing trajectories, bats always refrained from emitting echolocation calls for a short while after any event. This post-event pause was longer after drinking (153 ms) than after any other event (9-64 ms; Fig. 3 & Fig. S3, tables S3/S4).

### Call duration

Mean duration (± sd) of the 20% loudest type I and type II search calls was 2.0 (±0.3) ms and 2.6 (±0.4) ms, respectively. The call duration of approach and buzz calls during object approach ranged between 1-2.4 ms (Fig. 5). During drinking and drinking attempt trajectories, call duration during the approach exceptionally seemed to initially increase and then remained approximately constant (Fig. 5). In contrast, in all other tasks, call duration was longest at the beginning of the approach and then decreased throughout the approach (Fig. 5). Maximum call durations varied between 1.7 ms (landing) and 2.4 ms (capture); minimum durations at the end of the object approach were between 1.0 ms (prey captures) and 1.6 ms (drinking / drinking attempts; Fig. 5, tables S3/S4). As before, call duration dynamics were task-specific beyond those common temporal patterns (Fig. 5). Bats controlled call duration most dynamically when capturing prey, starting with the longest (2.4 ms) and ending with the shortest call duration (1.0 ms). Thus, they more than halved their call durations and, accordingly, the call-echo-overlap zone (Schnitzler & Kalko, 2001; Wilson & Moss, 2004), i.e., the range where returning echoes overlap in time with the outgoing call. Shortening of call duration started generally slow and was most pronounced just prior to prey captures. In contrast, landing approaches directly started with shorter call durations (1.7 ms) that also changed only slowly. Thus, call durations during landing were the shortest of all tasks throughout the approach, except for the last ∼70 ms, when call durations were shorter during capture (attempts). When exiting, shortening of call duration started the latest of all tasks, and then followed a similar linear trend as during capture. During drinking (attempts), bats used the longest call duration of all tasks at the end of a task (1.6 ms, Fig. 5, tables S3-S5).

**Fig 5:**
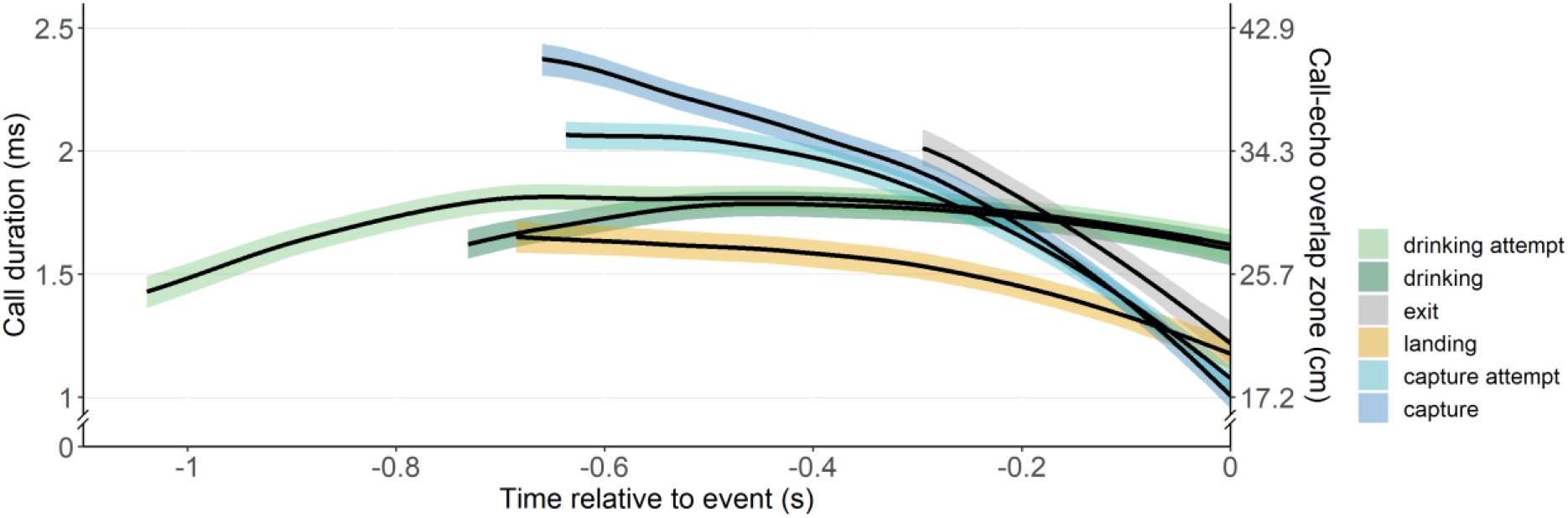
Dynamics of call duration (left y-axis) and call-echo overlap zone (right y-axis) during object approach. Call-echo overlap zone indicates the bat-object distance below which returning echoes overlap in time with the outgoing call; it was calculated assuming a speed of sound of 343 m/s. Depicted are model predictions (solid lines) and 95% credible intervals (coloured bands). The model is based on N=4322 calls. Predictions are made starting from the event-specific median start time of the final object approach phase.

### Frequency changes during object approach

Bats changed the spectral characteristics of their calls during object approach, with some task specific differences. These spectral changes were not so much achieved by shifting the peak frequency of each harmonic (Fig. 6a), but by changing energy distribution between harmonics (Fig. 6b). Mean peak frequencies of the first and second harmonics over the entire final object approach were rather constant at 39-43 kHz and 71-85 kHz, respectively. At the beginning of the object approach, almost all calls (91-99%) had peak frequencies within the first harmonic. Only landing trials started with a substantial proportion of calls (26%) having their peak frequency in the second harmonic (Fig. 6b), which then further increased earlier and more pronounced than during drinking, exit, and prey captures (Fig. 6b). Interestingly, bats used a higher proportion of calls with peak frequencies in the second harmonic during successful than unsuccessful prey capture trials (Fig. 6b, see non-overlapping confidence intervals; Table S5). Towards the end of the object approach, the proportion of calls with peak frequency in the second harmonic went back to pre-approach levels in most tasks. Only during drinking trials, this proportion started to increase later and kept increasing until the very end of the object approach. In consequence, at the end of the object approach, the proportion of calls with peak frequency in the second harmonic was substantially higher during drinking trials than during any other task (Fig. 6b, table S5).

**Fig 6:**
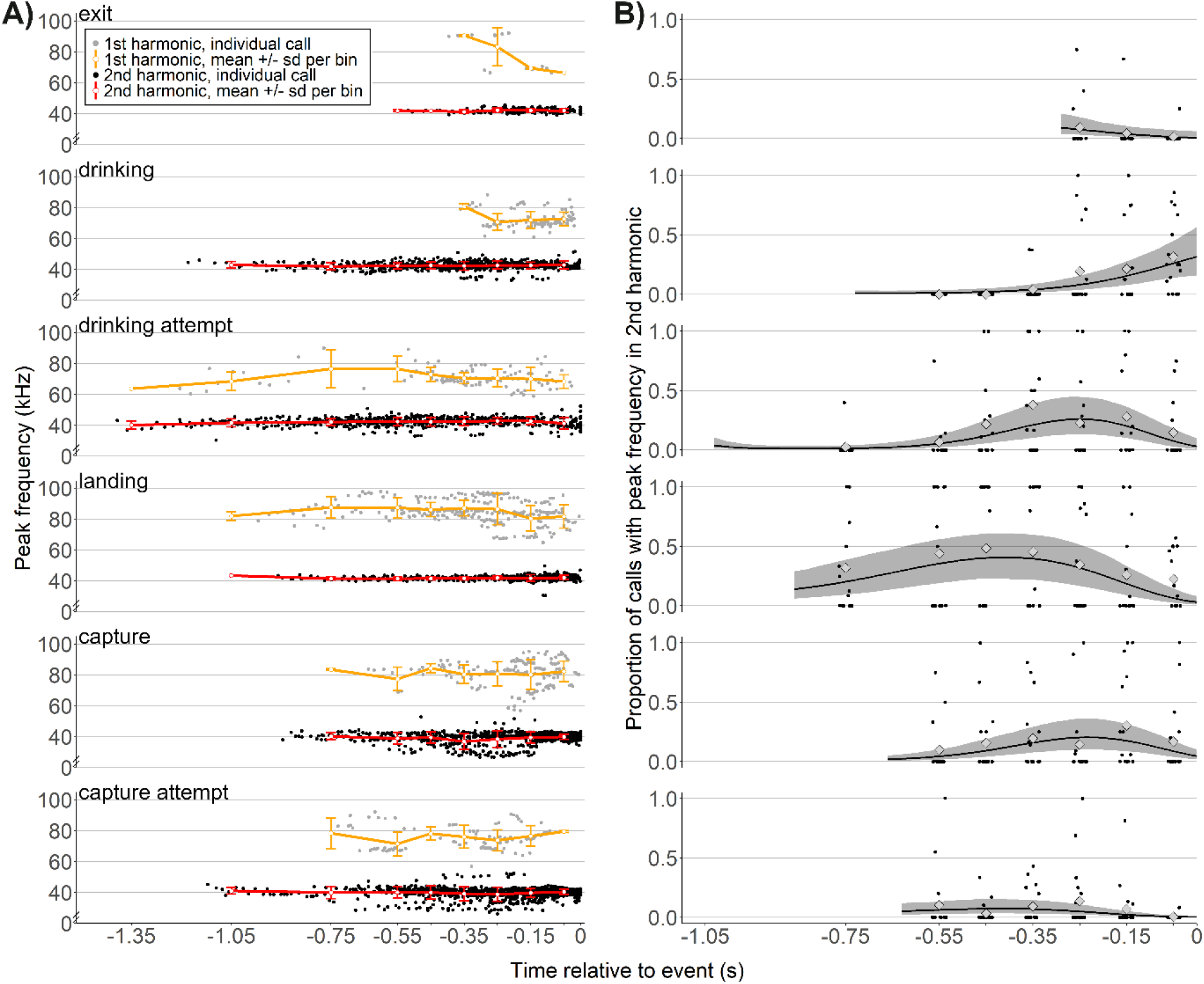
Dynamics of call peak frequency during object approach. A) Peak frequency of the first (red) and second (orange) harmonics did not change substantially over the course of the final object approach. Dots: individual calls; coloured lines: mean ± sd per time bin, i.e. per 0.3 s or per 0.1 s for calls emitted before or after -0.6 s relative to the event, respectively. B) Proportion of calls with most energy in the second harmonic, i.e. at higher frequencies. Depicted are posterior mean predictions (solid lines) and credible intervals (grey areas) from a generalised mixed model with binomial error distribution. We also show the underlying data, i.e. the proportion of calls with peak frequency in the second harmonic per trajectory and time bin (black dots, jittered around bin centre, N=742). Further, for each time bin we overlaid the mean proportion of calls with peak frequency in the second harmonic (light grey diamonds, calculated over all trajectories of any given task), which fit well the model predictions.

## Discussion

Here, we demonstrate that the aerial-hawking insectivorous barbastelle bat *Barbastella barbastellus* adjusts various aspects of its active sensory sampling strategy in a task-dependent manner. We show that barbastelle bats start their sensory target approach behaviour later and increase call rate less when approaching an opening than when drinking, landing, or capturing prey. On the contrary, bats only emitted buzz sequences with highest information update rates and shortest call durations when attacking prey. Lastly, bats used high information update rates and high sound frequencies for high spatial accuracy the most when landing, i.e., approaching a solid surface. Thus, we argue that the echolocation adjustments the bats made during target approach reflect the sensory-motor difficulty of any given task. Our results suggest that it is comparably easy for bats to navigate through an opening but challenging to capture small prey, and that bats adjust their echolocation parameters in a task-specific way matching its difficulty. Thus, in the following we discuss the challenges bats face in a task-specific manner before widening the scope to other taxa and discussing the lessons learned from comparing across tasks.

### Navigating through an opening

Presumably, among the tasks we studied it is easiest for echolocating bats to exit through an opening, particularly since the opening we offered was comparably large (0.5 x 1.3 m, height x width; wingspan of barbastelles is 26-30 cm (Skiba, 2009)), constant and predictable. As long as bats reliably headed towards the opening inside the wall and not the wall *per se*, ranging did not need to be particularly precise. Additionally, the bats’ spatial memory (Barchi, Knowles, & Simmons, 2013; Thiele & Winter, 2005) likely reduced the need for high update rates before exiting, particularly since bats had been used to the wall, the opening and surrounding landmarks in the flight room for about three weeks before we started to record exit trajectories. Indeed, big brown bats *Eptesicus fuscus* use landmarks to navigate an opening in a net (Jensen, Moss, & Surlykke, 2005). They also emit longer call intervals when flying through a larger compared to a smaller opening (Falk et al., 2015), suggesting that the need for high update rates decreases with increasing size of an opening; and the opening in our study was even larger than the one used by Falk et al. (2015). We thus suggest that the type of sensory-motor challenge and the bats’ experience and memory rendered the exit task the least challenging one. Accordingly, compared to the other three tasks, bats initiated their approach the latest and used only few approach calls with low update rates when navigating through the opening. Arguably, flight speed during exit is faster than during the other tasks. Together with the long call intervals during exit, this means that the distances flown between subsequent emissions are even larger during exit than in all other tasks, reemphasising that exiting through an opening poses the least sensory-motor challenges of all tasks tested. The larger flight speed and later onset of approach could potentially mean that the bats use distance to object to initiate their approach (in contrast to time to contact that we analysed). However, previous three-dimensional tracking of barbastelle prey capture behaviour under two different conditions showed no such pattern: when attacking tethered prey in the field and flight room, barbastelles flew at different speeds (7.4 vs. 3.1 m/s) and started their approach at different distances (1.6 vs. 1.2 m) and times to contact (200 vs. 390 ms) (Lewanzik & Goerlitz, 2018). This shows that the bats adjust their sensory-motor behaviour flexibly not only to the actual task, but also to various aspects of the environment, such as its spatial layout, its sound reflectivity and the available space for flight manoeuvres.

Interestingly, bats substantially increased between-group intervals and even emitted a few search calls already before passing the opening, indicating that they already shifted their sonar gaze to positions behind the opening. Despite this increase in call interval to allow for echoes arriving from beyond the wall, the bats continued to reduce their call duration to ∼1.2 ms until about 20 cm distance to the wall, thereby avoiding temporal overlap between calls and echoes from the close wall (call-echo ambiguity; Beetz, Kössl, & Hechavarría, 2019; Moss, Chiu, & Surlykke, 2011). Thus, even in this rather simple task, without a subsequent mealworm to catch (Surlykke et al., 2009) or a second hole to navigate through (Falk et al., 2015), the bats used a sensory strategy for simultaneous sampling of short- and long-range objects.

### Drinking

Bats innately recognise water bodies due to their mirror-like echo reflection properties (Greif & Siemers, 2010). To drink on the wing, bats must get low enough to reach the water with their tongue while avoiding touching the water surface with wings or body. Simultaneously, they must keep track of the scene ahead and plan their trajectory. Hence, drinking on the wing likely poses more sensory-motor challenges than flying through a large opening. Indeed, bats initiated their approach much earlier and called more frequently when approaching the pond compared to the opening. At the same time, drinking is likely less challenging than attacking prey since the position of any water surface is more predictable than that of a small and unpredictably moving prey. Indeed, bats reduced call intervals less when drinking than when landing or capturing prey, and never emitted buzz calls (<6ms inter-call-interval), corroborating previous findings of less pronounced increases in call rate during drinking than prey capture (Russo *et al*. 2016). In addition, call intervals during drinking need to be long enough to allow echo returns from distant objects before emitting the next call, in order for the bat to plan its prospective flight trajectory. With a minimum call interval of 10.7 ms during drinking approaches, bats could receive echoes from about 1.8 m distance without call-echo ambiguity, indicating that bats likely kept track of the edge of the pool to avoid collisions.

When flying at close distance above water, echoes of the water surface vertically below the bat arrive with very short delay (e.g. <0.3 ms at 5 cm above water). Most likely, bats will have difficulties to emit calls that are short enough to avoid call-echo overlap at all under such close-range conditions, especially given that the middle-ear reflex prevents echo detection shortly after call emission (Henson Jr., 1965; Kick & Simmons, 1984; Suga & Schlegel, 1972). This would explain why barbastelles ended their approach with longer call durations when approaching the pond compared to all other tasks. Our behavioural data thus match a spectral interference-based sensory strategy for estimating flight height above water surfaces (Hoffmann et al., 2015): as the outgoing call and returning echo overlap in time when flying close above the water surface, the flight height determines position and magnitude of spectral notches generated through interference between call and echo. Thus, when drinking, bats probably benefit from emitting comparably long calls that allow for sufficient temporal overlap between call and echo. Our observed initial increase in call duration, however, is likely a measurement artefact, as calls emitted at large distance had comparably low received amplitudes and thus relatively short call duration measurements.

The pause between the event and the first post-event call was largest after drinking (150 ms), corroborating long post-drinking pauses (200-350 ms) in other species (Kloepper, Simmons, & Simmons, 2019). Presumably, bats first need to swallow the water before being able to exhale and open the mouth for emitting the next call. In contrast, bats might well emit these calls while holding a prey item between their teeth. This conclusion is supported by shorter post-event pause (∼50 ms) after capture and the observation that barbastelles usually eat their prey in flight, which – depending on prey size – can take considerable time. Interestingly, the first call(s) after capture and drinking events where type II calls that presumably are emitted nasally – which could suggest that nasal emission is easier than mouth emission while and after chewing and swallowing. However, bats also emitted type II calls after landing, which could either indicate another reason for preferring nasal over mouth emission while being stationary, or yet another reason common to all tasks.

### Landing

Bats need to execute a complex, almost acrobatic aerial rotation to land head-under-heels (Riskin et al., 2009). Probably, the major challenge for bats landing on a wall is to execute that manoeuvre at the right position and right time, such that they are correctly positioned once reaching the wall. Thus, regular and precise ranging is of paramount importance prior to landing. Accordingly, the information update rate was higher when approaching the landing site than when approaching the drinking site. Similarly, both tongue-clicking fruit bats (Danilovich et al., 2015) and laryngeally echolocating bats (Hiryu, Hagino, Riquimaroux, & Watanabe, 2007b; Melcón, Denzinger, & Schnitzler, 2007; Russo et al., 2007) increased their call rates just before landing, suggesting that high information update rates are a general necessity for landing in animals moving fast in 3D space. Likewise, call durations during landing were the shortest of all tasks for most part of the approach, enabling precise distance information and little forward-masking early on during the approach (Boonman, Parsons, & Jones, 2003; Schnitzler & Kalko, 2001).

After successful landings, bats continued to call. Theoretically, this might be because we misidentified the time of landing, but we are confident that this was not the case, even if the wall contact was not visible in the video or the spectrogram: (i) the sound from hitting the wall was always audible in the sound file. (ii) Even when the sound was also visible in the spectrogram, and the timing thus less ambiguous, bats continued calling after landing. (iii) At the times defined as wall contact, within-group call intervals were shortest while between-group intervals were already increasing again (Fig. 4). In addition, also horseshoe bats continued calling after landing (Tian & Schnitzler, 1997). We assume that bats continue calling after wall contact for multiple reasons: (i) in case landing fails, continued sensory sampling would allow to evaluate possible ‘emergency’ flight trajectories. (ii) Bats often crawled to adjacent and more sheltered sites after landing and therefore sampled the specific surrounding of the landing spot. (iii) Lastly, bats also need to sense their general environment after landing.

### Capturing prey

Barbastelles started their approach on average 0.7 s before prey capture, which is just slightly longer than the 0.4 s we measured previously in the flight room with a stationary prey (Lewanzik & Goerlitz 2018) and similar to drinking and landing. Yet, only when capturing prey did the bats always emit a buzz sequence with shortest call intervals and durations. Such high sampling rates (up to >200 calls/s) prior to capture are ubiquitous among echolocators, including all aerial-hawking insectivorous bats and toothed whales (Fais, Johnson, Wilson, Aguilar Soto, & Madsen, 2016; Jensen, Keller, Tyack, & Visser, 2020; Madsen & Surlykke, 2013; Ratcliffe et al., 2013; Wisniewska et al., 2016). This suggests (i) that prey capture is a major sensorimotor challenge for all echolocating animals foraging on prey moving in three dimensions and (ii) that the terminal buzz is a key functional adaptation for these animals (Ratcliffe et al., 2013). The maximum call rate measured here, 286 calls/s, is among the highest measured to date for bats (Ratcliffe et al., 2013; Russo et al., 2016) and might be adaptive for hunting the barbastelle’s typical prey, eared moths (Goerlitz et al., 2010). In contrast to earless prey that cannot hear and react to attacking bats, eared moths try to escape attacking bats with evasive flight. The barbastelle’s main strategy for hunting eared moths is to emit low-intensity echolocation calls that allow it to remain undetected until close to the prey (Lewanzik & Goerlitz, 2018). Due to the bats’ reaction time, these high call rates are unlikely to guide capture during the final stages of the attack just prior to contact (Geberl, Brinkløv, Wiegrebe, & Surlykke, 2015; Melcón et al., 2007). However, the high call rates might aid in tracking prey that initiates evasive manoeuvres shortly before capture despite the low call levels, or for retrospectively evaluating a failed attack (Melcón et al., 2007). Additionally, high call rates improve the perception and evaluation of object movement (Goerlitz, Geberl & Wiegrebe 2010, Baier & Wiegebe 2018, Baier, Stelzer & Wiegrebe 2018), which would allow the barbastelle bat, despite being a low-duty cycle bat, to perceive and track the wing beat of its prey. This might enable barbastelle bats to resolve moving prey from a stationary background and even to estimate prey size based on wing beat frequency, for selecting large and more profitable moths. To date, this was only shown for high-duty cycle bats that emit very long and constant-frequency echolocation calls (Koselj, Schnitzler & Siemers 2011).

During unsuccessful prey capture attempts, and in contrast to successful prey captures, bats kept the peak frequency of almost all calls in the first harmonic throughout the final approach (Fig. 6b). This difference indicates that the bats either failed to capture prey because they did not control their sensory information appropriately, or they approached the prey using a suboptimal trajectory and hence purposely kept most energy at lower frequencies to have a broader sonar beam, i.e. a broader ‘field of view’, in such trials. If bats approached the prey in a way that put them at risk to collide with the stand, for instance, a broader beam could have allowed them to track both, prey and stand; yet focusing only partially on the prey might have been insufficient to successfully capture it.

### General discussion

By comparing the echolocation behaviour of barbastelle bats during object approach across different tasks, we here demonstrate that bats adjusted various call parameters independently from each other in a task- and difficulty-dependent manner – as they do to improve signal detectability in noise (Luo, Goerlitz, Brumm, & Wiegrebe, 2015). Our results imply that bats adjust the acquisition of both information quantity and quality independently to the sensory constraints of each task. Interestingly, they did so from the very beginning of the approach. This implies that the bats decide on a task-specific object approach behaviour based on information exclusively received from search call echoes. With the onset of approach echolocation, they already have decided about their sensory-motor program for object approach (Amichai & Yovel, 2017), which they then execute in a goal-directed and task-specific manner, focusing echolocation on the target (Koblitz et al., 2011; Surlykke, Ghose, & Moss, 2009).

Comparisons of onset of object approach between tasks might be affected by our measure of the reference (event) time. While we used the actual contact with the object as reference time for drinking and landing, we used the last recorded call and last buzz call for exit and capture, respectively. However, we argue that this difference is small and does not affect our conclusions. When exiting through the opening, the maximal temporal error between the last detected call and the actual passage of the window equals one call interval, i.e. maximally 44 ms (Fig. 4; conservatively assuming long between-group call intervals), but likely much less. For capture, the last buzz call coincides well with the time of prey contact in Daubenton’s bats *Myotis daubentonii* when aerial hawking (Geberl et al., 2015) and is emitted on average 10 ms before contact in Natterer’s bats *Myotis daubentonii* (Melcón et al., 2007). Thus, the potential bias introduced by using the last call and last buzz call instead of the actual time of contact is small compared to the effects we present in this paper.

Other echolocating animals, such as toothed whales, face very similar trade-offs for adjusting information quantity and quality. Despite the large differences in body size and physical conditions of their environments, all echolocating mammals resolve these trade-offs in strikingly similar ways (Madsen & Surlykke, 2013). For example, all echolocators need to emit signals frequently enough to avoid obstacles and to track prey, but suffer from echo ambiguity when the next signal is emitted before all echoes have returned. Both bats and toothed whales reduce the time interval between consecutive signals in such a way to maximise information flow while avoiding echo ambiguity when approaching objects (Jensen, Bejder, Wahlberg, & Madsen, 2009; Ladegaard, Jensen, Beedholm, da Silva, & Madsen, 2017; Melcón et al., 2007). Both taxa also emit a final buzz for fine-scale tracking prior to capture and reduce output intensity during the approach to limit detection range and to reduce the complexity of their auditory scene (Jensen et al., 2020; Ladegaard et al., 2017; Madsen & Surlykke, 2013). Thus, all echolocating animals actively control acoustic information flow and their sensory volume to guide subsequent actions. Here, we confirm that this active control is not connected in a general time- or distant-dependent manner to the approached object, but is finely adjusted in a task-specific way to four different tasks differing in sensory-motor difficulty.

Not only echolocators but every animal including humans needs to constantly acquire information about its environment to make appropriate motor decisions when moving about and interacting with others (Dall et al., 2005). Under some circumstances, social information provided by others might be exploited (Lewanzik et al., 2019; Valone & Templeton, 2002), but social information cannot completely substitute personal information. This study contributes to our understanding of how animals adjust the acquisition of personal information to deal with the constraints they face when sensing the world sequentially through sound. Our results show that they increase information quantity (call rate) only to a level as required by the task but not above that, likely to minimize the costs required for vocal production (Elemans et al., 2011; Currie et al 2020) and sensory analysis (Laughlin et al., 1998; Niven & Laughlin, 2008) at high information update rates. In addition, by flexibly adjusting call features, such as duration and frequency, bats also adapt the quality of the acquired information. Hence, our study highlights the importance of behavioural flexibility to optimize the cost-benefit ratio of sensing and the quantity and quality of sensory information.

Apart from the strategies investigated here, actively sensing animals can further adjust information acquisition by controlling the beam shape of emitted energy. Narrower beams prevent uninformative side-echoes while broader beams widen the field of view. Investigating how changes in beam shape relate to changes in other signal parameters would further our understanding of how animals interact with their environment. Here, barbastelle bats might be particularly interesting subjects, given their alternating emission through mouth and nose and the absence of a substantial drop in frequency at the very end of the approach which most other vespertilionid bats use to broaden their beams. Do barbastelle bats broaden their beams by other means than lowering frequency, e.g. by changing emission aperture (mouth opening, relative nostril position)? Also, simultaneously tracking flight trajectories while recoding call parameters and beam shape during various tasks would add another level of information. Given that barbastelles most likely emit their approach calls upwards through the nostrils, do they have to approach targets from below? Do flight and echolocation behaviour condition each other? Many exciting avenues are still to be explored.

## Acknowledgements

We thank the authorities for granting capture and experimental permits (55.1-8646.1-1 and 55.2-1-54-2532-18-2015, respectively) and the Max Planck Institute for Ornithology for providing facilities and logistic support. Particularly, we thank Christian Völk and Felix Hartl for valuable logistic support. Further, we are grateful to all members of the Acoustic and Functional Ecology Group and to an anonymous reviewer for constructive feedback on earlier versions of the manuscript.

## Authors’ contributions

H.R.G. and D.L. conceived the idea and designed methodology; D.L. collected and analysed the data and led the writing of the manuscript. All authors contributed critically to the drafts and gave final approval for publication.

## Data availability statement

All data is available as supplementary information on the journal website.

