## Supplemental text, figures and tables for "Task-dependent vocal adjustments to optimize biosonar-based information acquisition"

### SUPPLEMENTARY INFORMATION

### Supplementary methods, results and references

#### Effect of the "flutter simulator" (propeller) to simulate moth wingbeats:

**Methods:** To test if the flutter simulator that simulated moth wingbeats influenced any of our response variables in the prey capture trials, we fit generalised linear mixed-effects models to the response variables using only data from prey capture trials. We fit models with a Gaussian error distribution to our response variables 'onset of final approach' (rel. to the event time), 'post event pause', 'call duration', and 'inter-call interval' (both within-group and between-group) using the lmer function of the lme4 package (Bates, Maechler, Bolker, & Walker, 2015). 'Onset of final object approach' and 'post event pause' were modelled as a function of the fixed effects 'event type' (categorical with two levels: prey capture, and unsuccessful prey capture attempt) and 'flutter simulator presence' (categorical with two levels: yes / no) and – to account for repeated measures – the random effect 'bat ID' (categorical with three levels). 'Call duration' and 'inter-call interval' were modelled as a function of the fixed effects 'time before event' (continuous), 'event type', and the interaction between the two. Further, we included a 2$^{nd}$ order polynomial of 'time before event' as well as its interactions with 'event type' as predictors in the models since 'time before event' had non-linear effects on these response variables. To prevent autocorrelation, we also included 'call duration of previous call' or 'call interval of previous call' (continuous) as predictor in these models. Random effects included were 'trajectory no' (categorical with 35 levels) to account for potential dependency among calls within the same trajectory and 'bat ID'.

To model changes in the proportion of calls with peak frequency in the 2$^{nd}$ harmonic over time, we first calculated the number of calls per 0.3 s time bins with peak frequency in the 1$^{st}$ and 2$^{nd}$ harmonic, respectively (i.e. < or >= 55 kHz; compare Fig. S2). We then fitted a generalised linear mixed model with binomial error distribution to the response variable 'proportion of calls with peak frequency in 2nd harmonic' (weighed by the total number of calls per bin) using the glmer function of the lme4 package; including the fixed effects 'time bin' (modelled as continuous variable) as a 3$^{rd}$ order polynomial, 'event type', the interaction thereof, and the random effects 'trajectory no' and 'bat ID'. In contrast to all other models, we here included data of the entire object approach call sequence (i.e. including also search and approach calls before the beginning of the final approach phase) because binning the data substantially reduced sample size and would not allow modelling polynomial effects otherwise.

All continuous predictors except 'time bin' were scaled by their standard deviation and (with the exception of 'time before event' and 'time bin') centred on their mean. Model assumptions were verified by plotting residuals versus fitted values and by inspecting QQ-plots of the model residuals and of the random effects. We assessed residuals for temporal dependency by plotting estimates of the autocorrelation function using R's acf function.

**Results:** The propeller did not significantly affect any response variable (Tables S1 & S2).

**References:** Bates, D., Maechler, M., Bolker, B., & Walker, S. (2015). Fitting linear mixed-effects models using lme4. *Journal of Statistical Software*, *67*, 1–48.

### Supplementary Figures

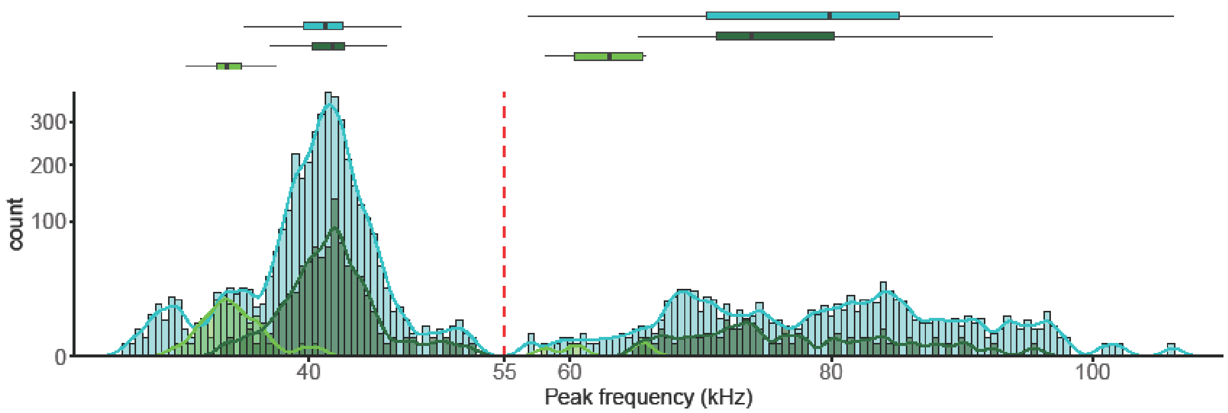


**Fig. S1**: Density plots overlaid on histograms of peak frequencies of type I search calls (light green), type II search calls (dark green), and approach calls (light blue). First and second harmonics were clearly separated by their peak frequency below/above 55 kHz (red dotted line). Distributions of peak frequencies call type and harmonic are depicted as horizontal box plots, showing median, quartiles, and largest value up to 1.5 * inter-quartile-range beyond the quartiles.

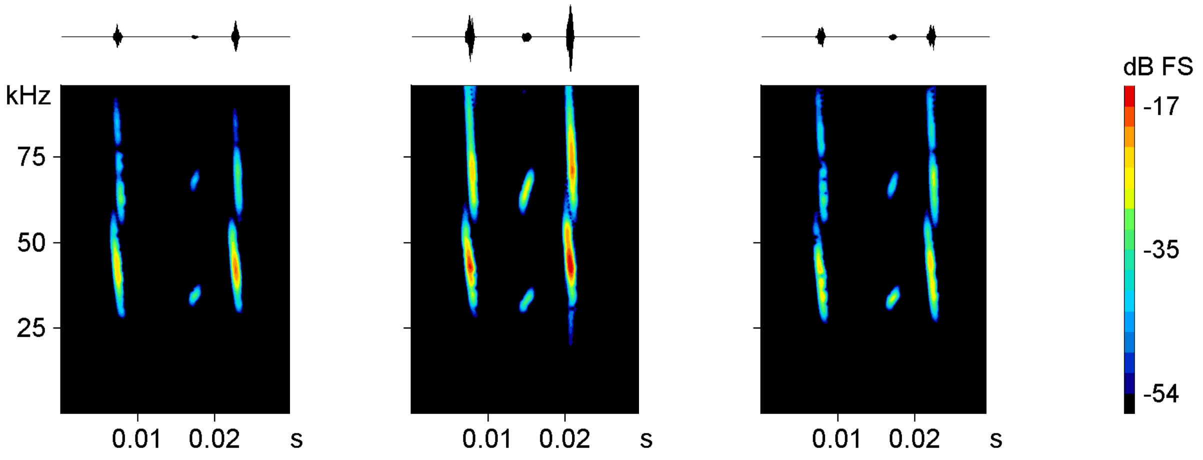


**Fig. S2**: Spectrograms (bottom) and oscillogram (top) of three UM-type (upward-modulated) calls (the faint central call in each spectrogram), each one surrounded by normal echolocation calls. UM-type calls were emitted by the same individual bat during the final approach before drinking or attempting to drink. We speculate that these UM-type calls were unintentionally emitted when the bat widely opened the mouth in preparation to drink and – maybe in consequence - did not precisely control the timing of exhalation and sound production, respectively.

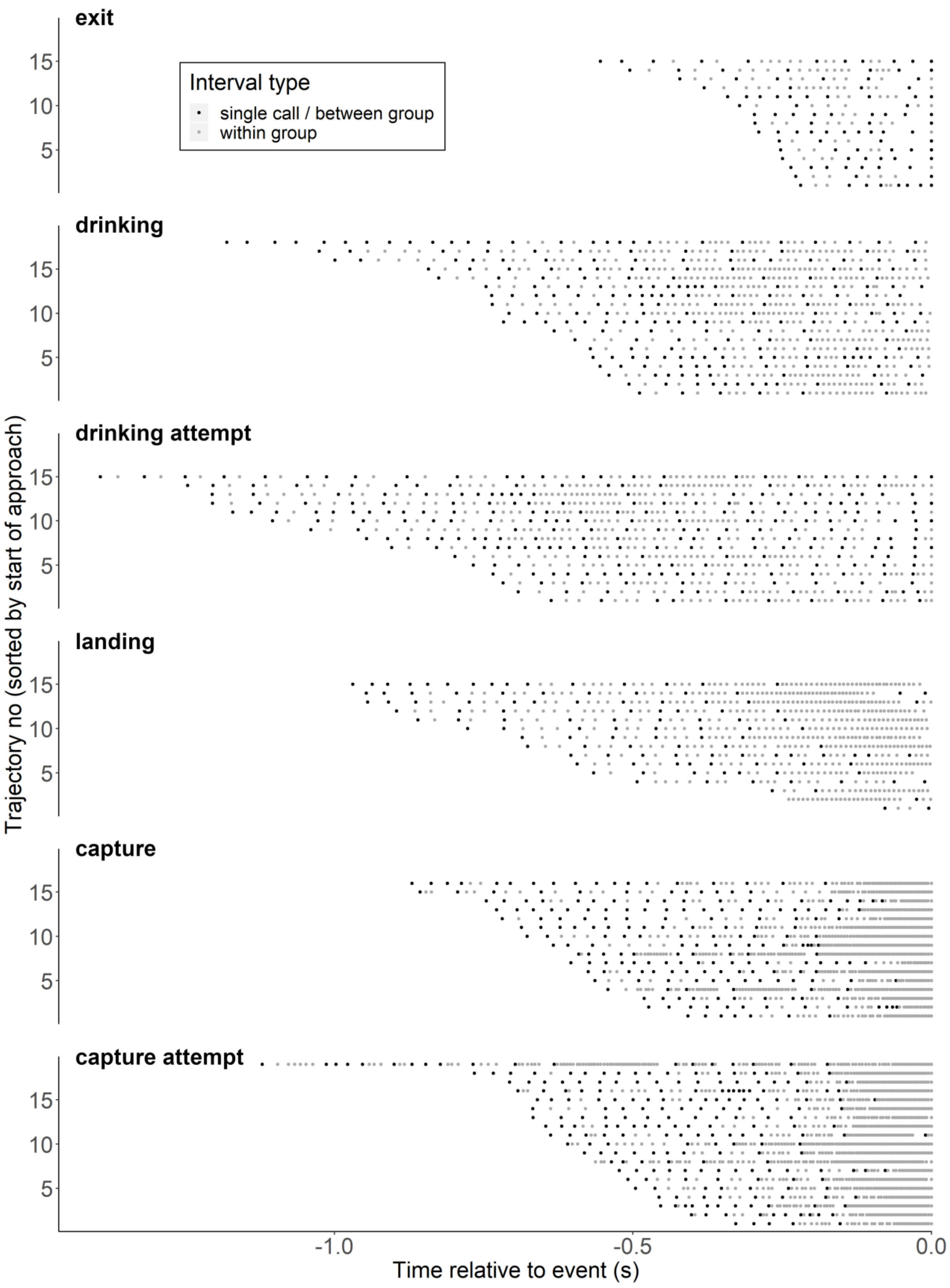


**Fig. S3**: Call grouping during the final object approach. Each call is grey-coded, indicating whether or not the interval to its preceding call is a within-group interval; if it is not a within-group interval, it can either be the first call of a call group or a single (ungrouped) call.

### Supplementary Tables

**Table S1**: Posterior mean estimates and 95% credible intervals (CIs) for fixed effects and interactions (indicated by ':') in models for seven measures of bats' acoustic flexibility when attacking prey (i.e. using data of successful captures and unsuccessful capture attempts only). Effects for which CI does not overlap with zero are indicated by asterisks. CIs for presence of our flutter simulator are highlighted in bold and overlap with zero in every model. Thus, the presence of the flutter simulator did most likely not affect any response variable.

| Response | Fixed effect or interaction | Estimate | Std. Error | T-value (z-value for 'binomial model') | 2.5% CI | 97.5% CI | |
|---|---|---|---|---|---|---|---|
| Start of final object approach | Intercept | -0,15 | 0,03 | -4,84 | -0,21 | -0,09 | * |
| | capture attempt | -0,02 | 0,02 | -0,69 | -0,06 | 0,03 | |
| | presence of flutter simulator | -0,02 | 0,02 | -0,93 | **-0,07** | **0,02** | |
| Within-group interval duration | Intercept | 5,56 | 0,21 | 26,73 | 5,15 | 5,98 | * |
| | time before event | -0,61 | 0,25 | -2,48 | -1,11 | -0,13 | * |
| | time before event^2 | 0,13 | 0,07 | 1,85 | -0,01 | 0,26 | |
| | capture attempt | -0,18 | 0,21 | -0,85 | -0,55 | 0,25 | |
| | previous call interval | 2,77 | 0,06 | 43,47 | 2,66 | 2,90 | * |
| | presence of flutter simulator | -0,10 | 0,14 | -0,71 | **-0,38** | **0,20** | |
| | time before event : capture attempt | -0,45 | 0,28 | -1,58 | -1,02 | 0,11 | |
| | time before event^2 : capture attempt | -0,24 | 0,08 | -3,16 | -0,38 | -0,09 | * |
| Between-group interval duration | Intercept | 12,66 | 2,52 | 5,02 | 7,67 | 17,28 | * |
| | time before event | -10,44 | 1,98 | -5,28 | -14,24 | -6,84 | * |
| | time before event^2 | -1,55 | 0,41 | -3,77 | -2,33 | -0,80 | * |
| | capture attempt | 1,91 | 3,06 | 0,62 | -3,84 | 8,24 | |
| | previous call interval | 1,26 | 0,37 | 3,40 | 0,54 | 1,94 | * |
| | presence of flutter simulator | -0,13 | 1,10 | -0,12 | **-2,37** | **2,09** | |
| | time before event : capture attempt | 2,53 | 2,51 | 1,01 | -2,16 | 7,52 | |
| | time before event^2 : capture attempt | 0,45 | 0,50 | 0,89 | -0,49 | 1,40 | |
| Start of final buzz | Intercept | -0,15 | 0,03 | -4,84 | -0,21 | -0,09 | * |
| | capture attempt | -0,02 | 0,02 | -0,69 | -0,06 | 0,03 | |
| | presence of flutter simulator | -0,02 | 0,02 | -0,93 | **-0,07** | **0,02** | |
| Post-event pause duration | Intercept | 0,04 | 0,01 | 3,80 | 0,02 | 0,07 | * |
| | capture attempt | 0,01 | 0,01 | 1,22 | -0,01 | 0,03 | |
| | presence of flutter simulator | 0,01 | 0,01 | 1,40 | **-0,01** | **0,03** | |
| Call duration | Intercept | 1,40 | 0,03 | 40,28 | 1,34 | 1,47 | * |
| | time before event | -0,25 | 0,03 | -9,61 | -0,29 | -0,19 | * |
| | time before event^2 | -0,04 | 0,01 | -6,29 | -0,06 | -0,03 | * |
| | capture attempt | 0,03 | 0,03 | 1,11 | -0,02 | 0,09 | |
| | duration of previous call | 0,45 | 0,01 | 57,15 | 0,43 | 0,46 | * |
| | presence of flutter simulator | 0,03 | 0,02 | 1,10 | **-0,02** | **0,07** | |
| | time before event : capture attempt | 0,07 | 0,03 | 2,43 | 0,02 | 0,11 | * |
| | time before event^2 : capture attempt | 0,02 | 0,01 | 2,15 | 0,00 | 0,03 | * |
| Proportion of second-harmonic calls | Intercept | -2,87 | 0,57 | -5,06 | 0,00 | -4,09 | |
| | capture attempt | -2,93 | 0,79 | -3,72 | 0,00 | -4,45 | |
| | time before event (binned) | -15,75 | 3,25 | -4,85 | 0,00 | -22,64 | |
| | time before event (binned)^2 | -42,66 | 8,90 | -4,80 | 0,00 | -61,19 | |
| | time before event (binned)^3 | -25,21 | 6,24 | -4,04 | 0,00 | -38,26 | |
| | presence of flutter simulator | -0,37 | 0,57 | -0,65 | **0,51** | **-1,41** | |
| | capture attempt : time before event (binned) | -3,66 | 5,05 | -0,73 | 0,47 | -13,18 | |
| | capture attempt : time before event (binned)$^2$ | 7,98 | 12,31 | 0,65 | 0,52 | -15,16 | |
| | capture attempt : time before event (binned)$^3$ | 7,01 | 8,12 | 0,86 | 0,39 | -8,33 | |

**Table S2**: Random effect variances and confidence intervals thereof from models for seven measures of bats' acoustic flexibility when attacking prey (i.e. using data of successful captures and unsuccessful capture attempts only).

| Response | Random effect | Variance | 2.5% CI | 97.5% CI |
|---|---|---|---|---|
| Start of final object approach | bat individual | 0,009 | 0,002 | 0,022 |
| Within-group interval duration | approach trajectory | 0,082 | 0,050 | 0,123 |
| | bat individual | 0,044 | 0,002 | 0,139 |
| Between-group interval duration | approach trajectory | 6,809 | 4,323 | 9,994 |
| | bat individual | 3,311 | 0,135 | 10,612 |
| Start of final buzz | bat individual | 0,002 | <0.001 | 0,004 |
| Post-event pause duration | bat individual | <0,001 | <0,001 | <0,001 |
| Call duration | approach trajectory | 0,005 | 0,003 | 0,006 |
| | bat individual | 0,002 | <0,001 | 0,005 |
| Proportion of second-harmonic calls | approach trajectory | 2,472 | 1,680 | 3,504 |
| | bat individual | <0,001 | <0,001 | <0,001 |

**Table S3**: Posterior mean estimates and 95% credible intervals (CIs) for fixed effects and interactions (indicated by ':') in models for seven measures of bats' acoustic flexibility during object approach. Effects for which CI does not overlap with zero are indicated by asterisks

| Response | Fixed effect or interaction | Estimate | Std. Error | T-value (z-value for 'binomial model') | 2.5% CI | 97.5% CI | |
|---|---|---|---|---|---|---|---|
| Start of final object approach | Intercept | -0,73 | 0,06 | -12,87 | -0,84 | -0,62 | * |
| | drinking attempt | -0,24 | 0,07 | -3,69 | -0,38 | -0,12 | * |
| | exit | 0,41 | 0,07 | 6,16 | 0,27 | 0,53 | * |
| | landing | 0,11 | 0,07 | 1,67 | -0,02 | 0,24 | |
| | capture | 0,09 | 0,06 | 1,32 | -0,04 | 0,21 | |
| | capture attempt | 0,12 | 0,06 | 1,95 | 0,00 | 0,23 | |
| Within-group interval duration | Intercept | 11,33 | 0,35 | 31,97 | 10,65 | 12,06 | * |
| | time before event | 2,22 | 0,76 | 2,93 | 0,70 | 3,80 | * |
| | time before event^2 | 0,99 | 0,46 | 2,14 | 0,06 | 1,93 | * |
| | time before event^3 | 0,06 | 0,08 | 0,71 | -0,11 | 0,21 | |
| | drinking attempt | 1,92 | 0,53 | 3,59 | 0,80 | 3,00 | * |
| | exit | 6,36 | 1,18 | 5,38 | 4,05 | 8,70 | * |
| | landing | -1,73 | 0,48 | -3,57 | -2,66 | -0,84 | * |
| | capture | -3,43 | 0,42 | -8,18 | -4,21 | -2,59 | * |
| | capture attempt | -3,52 | 0,41 | -8,63 | -4,33 | -2,71 | * |
| | previous call interval | 4,22 | 0,07 | 59,95 | 4,09 | 4,36 | * |
| | time before event : drinking attempt | 0,51 | 0,96 | 0,53 | -1,48 | 2,37 | |
| | time before event : exit | 21,97 | 4,63 | 4,74 | 12,98 | 31,43 | * |
| | time before event : landing | -0,66 | 1,11 | -0,59 | -2,87 | 1,51 | |
| | time before event : capture | -2,52 | 1,00 | -2,52 | -4,63 | -0,65 | * |
| | time before event : capture attempt | -2,82 | 0,88 | -3,19 | -4,55 | -1,03 | * |
| | time before event^2 : drinking attempt | -0,22 | 0,53 | -0,41 | -1,29 | 0,91 | |
| | time before event^2 : exit | 24,12 | 5,63 | 4,28 | 13,02 | 35,49 | * |
| | time before event^2 : landing | 0,66 | 0,71 | 0,92 | -0,80 | 2,03 | |
| | time before event^2 : capture | -0,48 | 0,68 | -0,70 | -1,92 | 0,81 | |
| | time before event^2 : capture attempt | -0,72 | 0,54 | -1,35 | -1,78 | 0,36 | |
| | time before event^3 : drinking attempt | -0,01 | 0,08 | -0,16 | -0,18 | 0,17 | |
| | time before event^3 : exit | 7,24 | 1,99 | 3,64 | 3,30 | 11,39 | * |
| | time before event^3 : landing | 0,22 | 0,13 | 1,73 | -0,04 | 0,47 | |
| | time before event^3 : capture | 0,01 | 0,13 | 0,04 | -0,25 | 0,25 | |
| | time before event^3 : capture attempt | 0,01 | 0,09 | 0,13 | -0,16 | 0,19 | |

**Table S3** (continued)

| Response | Fixed effect or interaction | Estimate | Std. Error | T-value (z-value for 'binomial model') | 2.5% CI | 97.5% CI | |
|---|---|---|---|---|---|---|---|
| Between-group interval duration | Intercept | 30,66 | 2,09 | 14,65 | 26,33 | 34,40 | * |
| | time before event | 13,66 | 3,66 | 3,73 | 6,78 | 20,95 | * |
| | time before event^2 | 9,51 | 2,11 | 4,50 | 5,43 | 13,61 | * |
| | time before event^3 | 1,35 | 0,33 | 4,08 | 0,73 | 1,98 | * |
| | drinking attempt | 6,49 | 2,84 | 2,28 | 1,14 | 11,87 | * |
| | exit | 19,17 | 3,08 | 6,23 | 13,37 | 25,21 | * |
| | landing | 2,79 | 3,98 | 0,70 | -4,81 | 10,22 | |
| | capture | -24,30 | 4,81 | -5,06 | -34,02 | -15,00 | * |
| | capture attempt | -14,85 | 4,51 | -3,29 | -23,09 | -6,36 | * |
| | previous call interval | 1,88 | 0,28 | 6,68 | 1,37 | 2,43 | * |
| | time before event : drinking attempt | 1,84 | 4,56 | 0,40 | -7,01 | 11,31 | |
| | time before event : exit | 53,60 | 10,57 | 5,07 | 31,16 | 73,20 | * |
| | time before event : landing | 10,46 | 7,47 | 1,40 | -4,36 | 24,92 | |
| | time before event : capture | -44,53 | 9,47 | -4,70 | -63,60 | -25,52 | * |
| | time before event : capture attempt | -23,35 | 7,75 | -3,01 | -38,44 | -9,02 | * |
| | time before event^2 : drinking attempt | -3,35 | 2,51 | -1,34 | -8,00 | 1,85 | |
| | time before event^2 : exit | 63,49 | 13,95 | 4,55 | 35,08 | 90,04 | * |
| | time before event^2 : landing | 8,38 | 4,54 | 1,84 | -0,55 | 17,22 | |
| | time before event^2 : capture | -23,20 | 5,91 | -3,93 | -34,59 | -11,60 | * |
| | time before event^2 : capture attempt | -10,98 | 4,24 | -2,59 | -19,29 | -2,99 | * |
| | time before event^3 : drinking attempt | -0,79 | 0,38 | -2,07 | -1,52 | -0,03 | * |
| | time before event^3 : exit | 18,95 | 4,95 | 3,83 | 9,14 | 28,31 | * |
| | time before event^3 : landing | 1,66 | 0,78 | 2,13 | 0,15 | 3,12 | * |
| | time before event^3 : capture | -3,37 | 1,10 | -3,05 | -5,41 | -1,21 | * |
| | time before event^3 : capture attempt | -1,25 | 0,67 | -1,86 | -2,65 | 0,06 | |
| Start of final buzz | Intercept | -0,16 | 0,03 | -5,48 | -0,21 | -0,10 | * |
| | capture attempt | -0,02 | 0,02 | -0,66 | -0,06 | 0,03 | |
| Post-event pause duration | Intercept | 0,15 | 0,02 | 7,25 | 0,11 | 0,20 | * |
| | drinking attempt | -0,09 | 0,03 | -3,20 | -0,15 | -0,03 | * |
| | landing | -0,14 | 0,03 | -5,38 | -0,20 | -0,09 | * |
| | capture | -0,10 | 0,03 | -3,88 | -0,15 | -0,05 | * |
| | capture attempt | -0,09 | 0,02 | -3,58 | -0,14 | -0,04 | * |
| Call duration | Intercept | 1,62 | 0,03 | 52,63 | 1,56 | 1,68 | * |
| | time before event | -0,06 | 0,03 | -2,23 | -0,10 | -0,01 | * |
| | time before event^2 | -0,01 | 0,01 | -2,05 | -0,03 | 0,00 | * |
| | drinking attempt | -0,06 | 0,04 | -1,71 | -0,14 | 0,01 | |
| | exit | -0,10 | 0,05 | -2,06 | -0,19 | 0,00 | * |
| | landing | -0,15 | 0,04 | -4,16 | -0,22 | -0,08 | * |
| | capture | -0,21 | 0,03 | -6,31 | -0,27 | -0,15 | * |
| | capture attempt | -0,17 | 0,03 | -5,54 | -0,24 | -0,11 | * |
| | duration of previous call | 0,36 | 0,01 | 71,43 | 0,35 | 0,37 | * |
| | time before event : drinking attempt | -0,06 | 0,03 | -2,03 | -0,13 | 0,00 | * |
| | time before event : exit | 0,00 | 0,10 | -0,03 | -0,20 | 0,19 | |
| | time before event : landing | -0,08 | 0,04 | -2,15 | -0,16 | -0,01 | * |
| | time before event : capture | -0,28 | 0,04 | -7,44 | -0,36 | -0,21 | * |
| | time before event : capture attempt | -0,19 | 0,03 | -5,90 | -0,26 | -0,13 | * |
| | time before event^2 : drinking attempt | -0,01 | 0,01 | -1,54 | -0,03 | 0,00 | |
| | time before event^2 : exit | 0,05 | 0,05 | 1,00 | -0,05 | 0,16 | |
| | time before event^2 : landing | -0,02 | 0,01 | -1,56 | -0,04 | 0,00 | |
| | time before event^2 : capture | -0,06 | 0,01 | -4,88 | -0,08 | -0,04 | * |
| | time before event^2 : capture attempt | -0,03 | 0,01 | -3,33 | -0,05 | -0,01 | * |

**Table S3** (continued)

| Response | Fixed effect or interaction | Estimate | Std. Error | T-value (z-value for 'binomial model') | 2.5% CI | 97.5% CI | |
|---|---|---|---|---|---|---|---|
| Proportion of second-harmonic calls | Intercept | -0,77 | 0,49 | -1,57 | -1,79 | 0,26 | |
| | drinking attempt | -2,65 | 0,78 | -3,41 | -4,17 | -1,16 | * |
| | exit | -4,18 | 1,10 | -3,79 | -6,47 | -1,82 | * |
| | landing | -2,69 | 0,71 | -3,79 | -4,09 | -1,32 | * |
| | capture | -2,25 | 0,66 | -3,39 | -3,63 | -0,94 | * |
| | capture attempt | -5,14 | 0,77 | -6,71 | -6,70 | -3,66 | * |
| | time before event (binned) | 5,03 | 3,33 | 1,51 | -2,57 | 11,81 | |
| | time before event (binned)^2 | -9,58 | 8,60 | -1,12 | -29,82 | 8,93 | |
| | time before event (binned)^3 | -12,51 | 5,66 | -2,21 | -25,72 | -0,40 | * |
| | drinking attempt : time before event (binned) | -26,20 | 5,16 | -5,08 | -36,36 | -15,36 | * |
| | exit : time before event (binned) | -19,96 | 7,14 | -2,80 | -35,44 | -4,26 | * |
| | landing : time before event (binned) | -22,36 | 4,44 | -5,04 | -31,32 | -13,20 | * |
| | capture : time before event (binned) | -20,68 | 4,56 | -4,53 | -29,85 | -10,19 | * |
| | capture attempt : time before event (binned) | -24,41 | 5,02 | -4,86 | -34,41 | -14,08 | * |
| | drinking attempt : time before event (binned)$^2$ | -45,56 | 12,51 | -3,64 | -71,69 | -18,23 | * |
| | exit : time before event (binned)$^2$ | -13,18 | 15,08 | -0,87 | -46,41 | 21,27 | |
| | landing : time before event (binned)$^2$ | -19,71 | 10,68 | -1,85 | -42,32 | 2,85 | |
| | capture : time before event (binned)$^2$ | -32,84 | 12,11 | -2,71 | -58,12 | -6,55 | * |
| | capture attempt : time before event (binned)$^2$ | -25,04 | 11,91 | -2,10 | -49,57 | -0,32 | * |
| | drinking attempt : time before event (binned)$^3$ | -21,32 | 7,97 | -2,67 | -38,71 | -4,13 | * |
| | exit : time before event (binned)$^3$ | 3,02 | 9,10 | 0,33 | -17,59 | 24,30 | |
| | landing : time before event (binned)$^3$ | -0,72 | 6,85 | -0,10 | -14,76 | 14,17 | |
| | capture : time before event (binned)$^3$ | -12,58 | 8,24 | -1,53 | -30,11 | 5,12 | |
| | capture attempt : time before event (binned)$^3$ | -5,67 | 7,57 | -0,75 | -21,70 | 10,57 | |

**Table S4**: Random effect variances and confidence intervals thereof from models for seven measures of bats' acoustic flexibility during object approach.

| Response | Random effect | Variance | 2.5% CI | 97.5% CI |
|---|---|---|---|---|
| Start of final object approach | bat individual | 0.004 | <0.001 | 0.009 |
| Within-group interval duration | approach trajectory | 0.106 | 0.079 | 0.138 |
| | bat individual | 0.003 | <0.001 | 0.010 |
| Between-group interval duration | approach trajectory | 25.535 | 21.281 | 30.465 |
| | bat individual | <0.001 | <0.001 | <0.001 |
| Start of final buzz | bat individual | 0.002 | <0.001 | 0.004 |
| Post-event pause duration | bat individual | <0.001 | <0.001 | 0.001 |
| Call duration | approach trajectory | 0.004 | 0.003 | 0.005 |
| | bat individual | 0.001 | <0.001 | 0.003 |
| Proportion of second-harmonic calls | approach trajectory | 2.010 | 1.676 | 2.409 |
| | bat individual | 0.029 | 0.002 | 0.084 |

**Table S5**: Post-hoc pairwise comparisons between all six levels of the fixed effect 'event type'. Contrasts were computed using the pairs function of the emmeans package in R for the time of the event (t = 0 s). P-values are tukey adjusted for multiple comparisons.

| Response | Contrast | Estimate | Std. Error | Df | T-ratio | P-value | |
|---|---|---|---|---|---|---|---|
| Start of final object approach | drinking - drinking attempt | 0.243 | 0.066 | 90.065 | 3.687 | 0.005 | ** |
| | drinking - exit | -0.406 | 0.066 | 90.065 | -6.155 | <0.001 | *** |
| | drinking - landing | -0.112 | 0.068 | 91.313 | -1.654 | 0.565 | |
| | drinking - prey capture | -0.086 | 0.065 | 90.098 | -1.319 | 0.774 | |
| | drinking - prey capture attempt | -0.121 | 0.062 | 90.298 | -1.943 | 0.383 | |
| | drinking attempt - exit | -0.649 | 0.069 | 90.000 | -9.433 | <0.001 | *** |
| | drinking attempt - landing | -0.355 | 0.070 | 90.962 | -5.072 | <0.001 | *** |
| | drinking attempt - prey capture | -0.329 | 0.068 | 90.027 | -4.851 | <0.001 | *** |
| | drinking attempt - prey capture attempt | -0.364 | 0.065 | 90.145 | -5.583 | <0.001 | *** |
| | exit - landing | 0.294 | 0.070 | 90.962 | 4.206 | <0.001 | *** |
| | exit - prey capture | 0.320 | 0.068 | 90.027 | 4.728 | <0.001 | *** |
| | exit - prey capture attempt | 0.285 | 0.065 | 90.145 | 4.368 | <0.001 | *** |
| | landing - prey capture | 0.026 | 0.069 | 91.175 | 0.378 | 0.999 | |
| | landing - prey capture attempt | -0.009 | 0.067 | 91.383 | -0.139 | 1.000 | |
| | prey capture - prey capture attempt | -0.035 | 0.064 | 90.054 | -0.554 | 0.994 | |
| Within-group interval duration (at time of event) | drinking - drinking attempt | -1.917 | 0.536 | 2032.728 | -3.578 | 0.005 | ** |
| | drinking - exit | -6.365 | 1.185 | 3092.231 | -5.371 | <0.001 | *** |
| | drinking - landing | 1.729 | 0.487 | 1775.698 | 3.547 | 0.005 | ** |
| | drinking - prey capture | 3.431 | 0.421 | 1487.238 | 8.144 | <0.001 | *** |
| | drinking - prey capture attempt | 3.520 | 0.411 | 1498.399 | 8.574 | <0.001 | *** |
| | drinking attempt - exit | -4.448 | 1.197 | 3078.421 | -3.716 | 0.003 | ** |
| | drinking attempt - landing | 3.645 | 0.528 | 1753.161 | 6.900 | <0.001 | *** |
| | drinking attempt - prey capture | 5.348 | 0.473 | 1491.558 | 11.306 | <0.001 | *** |
| | drinking attempt - prey capture attempt | 5.437 | 0.464 | 1509.402 | 11.706 | <0.001 | *** |
| | exit - landing | 8.093 | 1.180 | 3060.152 | 6.860 | <0.001 | *** |
| | exit - prey capture | 9.796 | 1.161 | 3054.713 | 8.436 | <0.001 | *** |
| | exit - prey capture attempt | 9.884 | 1.158 | 3059.388 | 8.535 | <0.001 | *** |
| | landing - prey capture | 1.702 | 0.398 | 1108.603 | 4.283 | <0.001 | *** |
| | landing - prey capture attempt | 1.791 | 0.384 | 1086.610 | 4.662 | <0.001 | *** |
| | prey capture - prey capture attempt | 0.089 | 0.276 | 508.116 | 0.321 | 1.000 | |
| Between-group interval duration (at time of event) | drinking - drinking attempt | -6.488 | 2.844 | 360.132 | -2.281 | 0.205 | |
| | drinking - exit | -19.171 | 3.081 | 447.593 | -6.223 | <0.001 | *** |
| | drinking - landing | -2.789 | 4.036 | 705.115 | -0.691 | 0.983 | |
| | drinking - prey capture | 24.303 | 4.810 | 927.245 | 5.052 | <0.001 | *** |
| | drinking - prey capture attempt | 14.854 | 4.510 | 937.473 | 3.294 | 0.013 | * |
| | drinking attempt - exit | -12.683 | 2.962 | 351.308 | -4.282 | <0.001 | *** |
| | drinking attempt - landing | 3.699 | 3.916 | 631.749 | 0.945 | 0.935 | |
| | drinking attempt - prey capture | 30.791 | 4.753 | 889.407 | 6.479 | <0.001 | *** |
| | drinking attempt - prey capture attempt | 21.342 | 4.442 | 894.410 | 4.804 | <0.001 | *** |
| | exit - landing | 16.381 | 4.086 | 680.912 | 4.009 | <0.001 | *** |
| | exit - prey capture | 43.474 | 4.901 | 907.489 | 8.870 | <0.001 | *** |
| | exit - prey capture attempt | 34.024 | 4.600 | 914.919 | 7.396 | <0.001 | *** |
| | landing - prey capture | 27.092 | 5.587 | 939.780 | 4.849 | <0.001 | *** |
| | landing - prey capture attempt | 17.643 | 5.307 | 953.876 | 3.325 | 0.012 | * |
| | prey capture - prey capture attempt | -9.450 | 5.882 | 1011.263 | -1.606 | 0.594 | |
| Post-event pause duration | drinking - drinking attempt | 0.093 | 0.029 | 72.610 | 3.180 | 0.018 | * |
| | drinking - landing | 0.144 | 0.027 | 73.757 | 5.285 | <0.001 | *** |
| | drinking - prey capture | 0.101 | 0.026 | 72.159 | 3.877 | 0.002 | ** |
| | drinking - prey capture attempt | 0.089 | 0.025 | 72.478 | 3.565 | 0.006 | ** |
| | drinking attempt - landing | 0.052 | 0.030 | 72.501 | 1.709 | 0.435 | |
| | drinking attempt - prey capture | 0.008 | 0.030 | 72.405 | 0.271 | 0.999 | |
| | drinking attempt - prey capture attempt | -0.004 | 0.029 | 72.560 | -0.125 | 1.000 | |
| | landing - prey capture | -0.043 | 0.028 | 73.624 | -1.558 | 0.529 | |
| | landing - prey capture attempt | -0.055 | 0.027 | 73.831 | -2.041 | 0.257 | |
| | prey capture - prey capture attempt | -0.012 | 0.026 | 72.088 | -0.455 | 0.991 | |

**Table S5** (continued)

| Response | Contrast | Estimate | Std. Error | Df | T-ratio | P-value | |
|---|---|---|---|---|---|---|---|
| Call duration (at time of event) | drinking - drinking attempt | 0,062 | 0,036 | 365,053 | 1,708 | 0,528 | |
| | drinking - exit | 0,098 | 0,048 | 953,298 | 2,058 | 0,311 | |
| | drinking - landing | 0,151 | 0,036 | 332,935 | 4,144 | <0.001 | *** |
| | drinking - prey capture | 0,208 | 0,033 | 265,977 | 6,303 | <0.001 | *** |
| | drinking - prey capture attempt | 0,174 | 0,031 | 260,255 | 5,539 | <0.001 | *** |
| | drinking attempt - exit | 0,036 | 0,049 | 893,456 | 0,737 | 0,977 | |
| | drinking attempt - landing | 0,089 | 0,038 | 334,817 | 2,339 | 0,182 | |
| | drinking attempt - prey capture | 0,146 | 0,035 | 271,622 | 4,192 | <0.001 | *** |
| | drinking attempt - prey capture attempt | 0,112 | 0,033 | 266,702 | 3,367 | 0,011 | * |
| | exit - landing | 0,053 | 0,049 | 831,160 | 1,080 | 0,889 | |
| | exit - prey capture | 0,110 | 0,046 | 770,202 | 2,382 | 0,164 | |
| | exit - prey capture attempt | 0,076 | 0,045 | 812,963 | 1,689 | 0,539 | |
| | landing - prey capture | 0,057 | 0,034 | 228,946 | 1,695 | 0,537 | |
| | landing - prey capture attempt | 0,024 | 0,032 | 224,825 | 0,727 | 0,978 | |
| | prey capture - prey capture attempt | -0,034 | 0,028 | 152,093 | -1,201 | 0,836 | |
| Proportion of second-harmonic calls (at time of event) | drinking - drinking attempt | 2,650 | 0,777 | NA | 3,411 | 0,009 | ** |
| | drinking - exit | 4,185 | 1,104 | NA | 3,790 | 0,002 | ** |
| | drinking - landing | 2,694 | 0,711 | NA | 3,787 | 0,002 | ** |
| | drinking - prey capture | 2,255 | 0,664 | NA | 3,395 | 0,009 | ** |
| | drinking - prey capture attempt | 5,145 | 0,766 | NA | 6,714 | <0.001 | *** |
| | drinking attempt - exit | 1,535 | 1,184 | NA | 1,296 | 0,787 | |
| | drinking attempt - landing | 0,043 | 0,818 | NA | 0,053 | 1,000 | |
| | drinking attempt - prey capture | -0,396 | 0,775 | NA | -0,510 | 0,996 | |
| | drinking attempt - prey capture attempt | 2,494 | 0,864 | NA | 2,888 | 0,045 | * |
| | exit - landing | -1,491 | 1,143 | NA | -1,305 | 0,782 | |
| | exit - prey capture | -1,930 | 1,111 | NA | -1,737 | 0,507 | |
| | exit - prey capture attempt | 0,960 | 1,174 | NA | 0,817 | 0,964 | |
| | landing - prey capture | -0,439 | 0,709 | NA | -0,619 | 0,990 | |
| | landing - prey capture attempt | 2,451 | 0,805 | NA | 3,046 | 0,028 | * |
| | prey capture - prey capture attempt | 2,890 | 0,759 | NA | 3,810 | 0,002 | ** |
